# Assessing scoring metrics for AlphaFold2 and AlphaFold3 protein complex predictions

**DOI:** 10.1101/2025.04.16.648930

**Authors:** Luca R. Genz, Sanjana Nair, Natan Nagar, Maya Topf

## Abstract

1.

Recent breakthroughs in AI-driven protein structure prediction have revolutionized structural biology, unlocking new possibilities to model complex biomolecular interactions. We evaluated widely-used scoring metrics for assessing models predicted by ColabFold with templates, ColabFold without templates, and AlphaFold3. We benchmarked the optimal cutoffs for these assessment scores using a set of 223 heterodimeric, high-resolution protein structures and their predictions. Our results show that ColabFold with templates and AlphaFold3 perform similarly and both outperform ColabFold without templates. However, the assessment scores perform best on ColabFold without templates. Furthermore, interface-specific scores are more reliable for evaluating protein complex predictions compared to the corresponding global scores. Notably, ipTM and *model confidence* achieve the best discrimination between correct and incorrect predictions. Based on our results, we developed a weighted combined score, C2Qscore, to improve model quality assessment. We used C2Qscore to analyse dimers from large assemblies solved by cryoEM, revealing potential limitations of the existing metrics when multiple configurations of heterodimers are possible. This study provides insights into the strengths and weaknesses of current scores and offers guidance for improving protein complex model assessment under realistic use case conditions. C2Qscore has been integrated as a tool into our ChimeraX plug-in PICKLUSTER v.2.0 and is also available as a command-line tool on https://gitlab.com/topf-lab/c2qscore.

**Impact of this work:** Many essential cellular functions rely on protein complexes, which are now predominantly predicted using AlphaFold by both experts and non-experts. This study systematically evaluates the performance of multiple widely used scoring metrics for distinguishing accurate from poor predictions. The new C2Qscore developed in this study improves the reliability of AlphaFold model assessments, enabling a more consistent and accessible evaluation of protein structures. These advancements support downstream applications in biomedicine, drug discovery, and computational protein design.

## 2. Introduction

Protein complexes are fundamental to various cellular processes, performing diverse functions in vivo (Schulz and Schirmer 2013). To understand the mechanisms of protein complex formation and macromolecular recognition, the precise identification and characterization of protein-protein interfaces is a crucial step (Xue et al. 2015). High-resolution structural data obtained from experimental methods such as X-ray crystallography and cryogenic-electron microscopy (cryo-EM) offer insights into these interfaces (Smyth and Martin 2000; Namba and Makino 2022). However, these are resource-intensive techniques and present challenges for large-scale structure determination (Xue et al. 2015). Recent advancements in AI-based protein structure prediction, particularly AlphaFold2, have mitigated these challenges by enabling large-scale structure predictions sometimes even with accuracies comparable to those of experimental methods (Jumper et al. 2021; Evans et al. 2022). This progress was recognized with the 2024 Nobel Prize in Chemistry. AlphaFold2’s successor, AlphaFold3, further advanced the field by allowing the prediction of protein complexes with lipids, nucleic acids and other ligands (Abramson et al. 2024).

Beyond predicting the structures of protein complexes, it is equally important to assess the accuracy of these predictions to ensure confidence in the models, which is crucial for downstream analyses, such as functional studies, protein engineering, and drug design (Baker and Sali 2001). However, evaluating the quality of predictions without a reference structure is challenging. The progress in the protein structure prediction field has led to the development of various methods for assessing model accuracy, with increasing emphasis on evaluating protein complexes (Studer et al. 2023 Oct 19). CASP (Critical Assessment of Structure Prediction), a community-wide experiment designed to track advancements in the structure prediction field, introduced the Estimation of Model Accuracy (EMA) category already in CASP7 to address this need (Cozzetto et al. 2007). To date, numerous prediction-based evaluation metrics have been proposed, most of which are machine learning-based (Studer et al. 2023 Oct 19). The graph neural network (GNN)-based scoring method, VoroIF-GNN (VoroIF) (Olechnovič and Venclovas 2023 Jul 21), emerged as one of the top performing scores in assessing interface quality in the CASP15 EMA iteration, ranking alongside ModFOLDdockR and ModFOLDdock (Edmunds et al. 2023 Jun 14) and VoroMQA (Olechnovič and Venclovas 2019). VoroIF uses Voronoi tessellation to derive interface graphs, providing a detailed, contact-based accuracy estimate for the entire interface. The VoroIF interface-specific score is calculated as the average score of interface residues (Average-VoroIF-GNN-residue-pCAD score), which is then adjusted for structural completeness based on the ratio of atoms in the model to atoms in the target.

Another commonly used metric for evaluating the quality of protein structure predictions is the predicted DockQ score (pDockQ) (Bryant et al. 2022). This score is derived by calculating the number of interfacial contacts and the average quality of interacting residues, which are then fitted to a sigmoid function of the DockQ score (Basu and Wallner 2016; Mirabello and Wallner 2024). Its more recent iteration, pDockQ2 (Zhu et al. 2023), was specifically developed for the assessment of multimeric protein complexes. To our knowledge, to date only Yin et al. (Yin et al. 2022) have conducted a systematic assessment of scores that evaluate AlphaFold2 multimeric predictions, using a relatively small benchmark set of 152 heterodimeric protein complexes. In their analysis, they compared the performance of AlphaFold2 to the protein-docking algorithm ZDOCK (Pierce et al. 2011). Evaluated metrics included the predicted Local Distance Difference Test (pLDDT) (Jumper et al. 2021), an interface-specific version of pLDDT, predicted Template Modeling score (pTM) (Jumper et al. 2021), an interface-specific Pairwise Alignment Error (PAE) (Evans et al. 2022), among other metrics, and these were compared against the interface Root Mean Square Deviation (I-RMSD) (Lensink and Wodak 2013; Basu and Wallner 2016). Furthermore, most of the current evaluation metrics, including pDockQ, VoroIF, and others, have been applied in isolation and were evaluated prior to the development of AlphaFold3.

In this study, we focused on the quality of the interface, by providing a comprehensive comparison and evaluation of commonly used scores for assessing predictions of heterodimeric protein complexes. We evaluated predictions from both AlphaFold2/ColabFold (Mirdita et al. 2022) (with and without templates) and AlphaFold3. We compared different scores, with DockQ as ground truth, including: interface predicted Local Distance Difference Test (ipLDDT) (Genz et al. 2023), pTM, interface pTM (ipTM) (Evans et al. 2022), *model confidence* (Evans et al. 2022), interface PAE (iPAE) (Evans et al. 2022; Genz et al. 2023), pDockQ2 (Zhu et al. 2023), and VoroIF. The individual scores were merged into one score, trained on predictions for 223 heterodimeric high-resolution structures from the Protein Data Bank (PDB) and tested on two independent datasets of predictions for (i) X-ray crystallographic structures and (ii) dimers from larger assemblies derived from cryoEM. The score has been integrated as a tool into our ChimeraX plug-in PICKLUSTER v.2.0 (Genz et al. 2023), providing users with interactive access to scoring metrics as a valuable analytical tool. This study highlights the strengths and limitations of current scoring methods and provides practical guidance for improving protein complex model evaluation in realistic use case scenarios.

## 3. Results

### 3.1. Benchmarking the prediction accuracy of ColabFold with templates, ColabFold template free and AlphaFold3

To evaluate the capability of ColabFold and AlphaFold3 in generating high-quality predictions of structural models, a comparative analysis was performed using a benchmark set. We focused exclusively on heterodimeric complexes rather than homodimeric ones, as AlphaFold2 generally performs better on homomeric interfaces (Evans et al. 2022; Zhu et al. 2023). By selecting heterodimers, we aimed to introduce greater diversity and a more challenging evaluation setting. Starting with 671 complexes, a filtering process (Methods, Suppl. Fig. 1) reduced the number of target structures to 257. Among these, 4 cases were identified in which the biological assembly (BA) assigned in the PDB is different from the asymmetric unit (AU), resulting in predictions resembling only one of the two evaluated using DockQ (Methods). This caused issues during alignment of the AU with the predicted model, leading to artificially low scores that do not reflect the prediction quality in these specific cases. To address this, only templates with the assigned dimeric BA identical to the AU were included in the benchmark set, resulting in a final set of 223 target structures.

Predictions were generated using ColabFold with templates (CF-T) and with the template-free (CF-F) option, and the AlphaFold3 web server (AF3). All predictions were performed with three recycles followed by relaxation (Hornak et al. 2006), producing five predictions per target structure. DockQ and individual prediction-based scores were calculated for a total of 1,115 models in each of the datasets-CF-T, CF-F, and AF3.

Using CAPRI (Critical Assessment of PRediction of Interactions) criteria for DockQ (“DockQ classification”, Methods) revealed that AF3 (39.8%) and CF-T (35.2%) had the highest proportion of ‘high’ quality models (DockQ > 0.8), making it the largest fraction for both prediction methods (Fig. 1A).

**Fig. 1.**
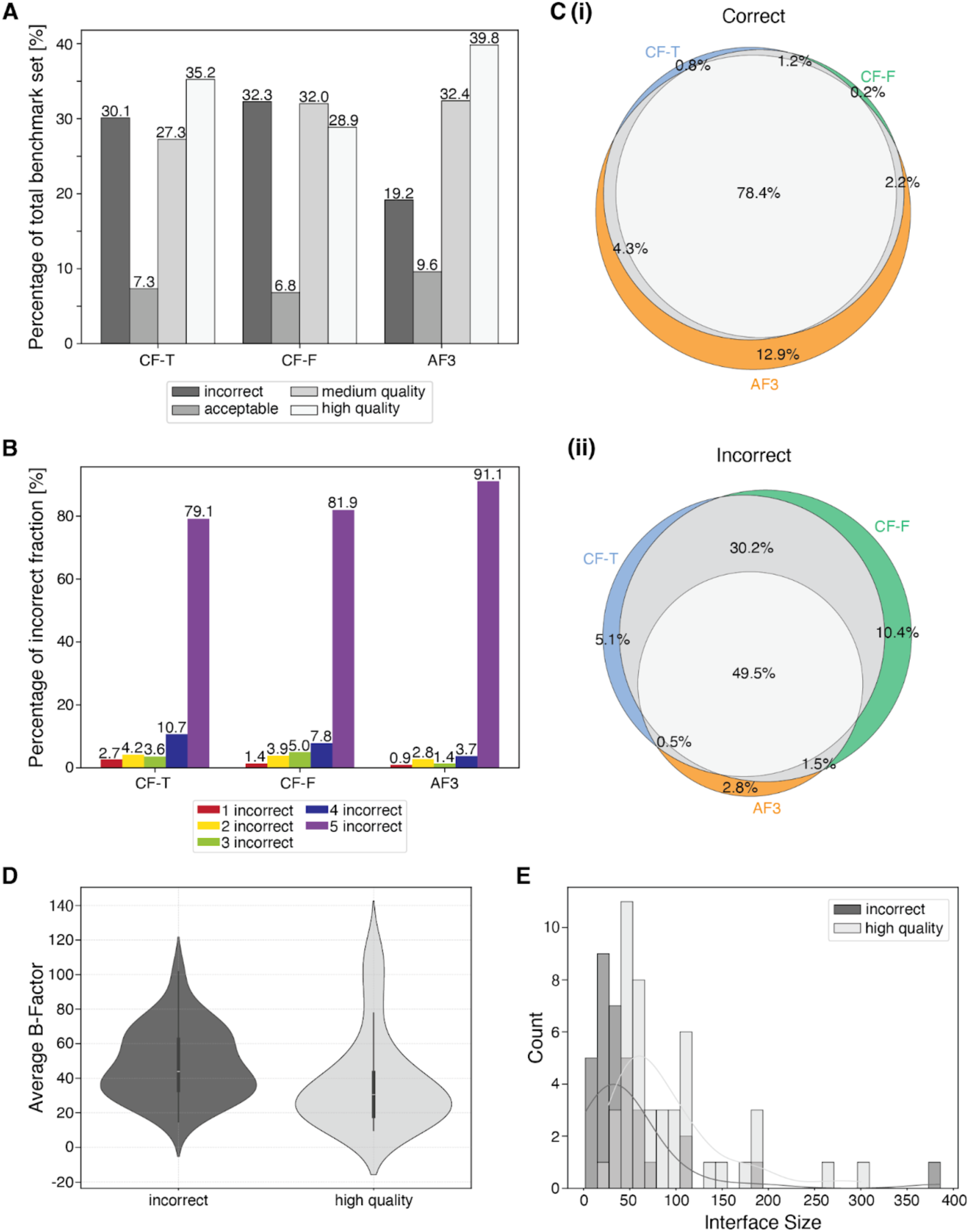
Quality distributions, overlap of DockQ classifications (5 levels) and detailed analysis of the targets across models predicted by CF-T, CF-F, and AF3. **A:** Quality assessment of the three sets based on DockQ classification criteria. **B:** Breakdown of the incorrect fraction from A, categorizing cases where all five models of a template are classified as incorrect, followed by cases with 4, 3, 2, and those with only one model classified as incorrect. **C(i):** Venn diagram of the ‘correct’ fraction (‘acceptable’, ‘medium’, ‘high’ are combined). The fraction of only CF-T predicting models of a specific quality are shown in blue, CF-F in green, and AF3 in orange. When two prediction methods agree, the fraction is represented in gray, and when all three prediction methods agree, it is shown in light gray. **(ii):** Venn diagram of the ‘incorrect’ fraction. Colors are as in (C(i)). **D:** Distribution of the average b-factor of the interface of targets that are consistently mispredicted vs. those that are consistently predicted with high quality across all three datasets. **E:** Interface size distributions of targets that are consistently mispredicted vs. those that are consistently predicted with high quality across all three datasets.

Of the three prediction methods, the CF-F set contained the lowest proportion of ‘high’ quality models at 28.9% with the largest fraction consisting of models of ‘medium’ quality. Analyzing the models classified as ‘incorrect’ using DockQ (DockQ < 0.23) showed that AF3 had the lowest percentage of incorrect models at 19.2%. In contrast, the CF-T and CF-F sets exhibited significantly higher proportions of incorrect models, at 30.1% and 32.3%, respectively.

Further investigation of the composition of the ‘incorrect’ fraction revealed that AF3 showed the highest percentage of cases where all five models of the prediction were classified as incorrect (91.1%) (Fig. 1B) CF-F revealed a significantly smaller proportion of cases where all five models of the prediction were classified as incorrect (81.9%) and the CF-T set had the lowest percentage of those (79.1%).

Considering only the ‘high’ quality fraction, AF3 predictions appear to be the most accurate. Moreover, if models classified as ‘acceptable’, ‘medium’ and ‘high’ quality according to the CAPRI criteria are all considered correct (as opposed to the ‘incorrect’ fraction), then AF3 predictions are superior, as they exhibit the lowest number of incorrect models. This detailed analysis of the quality of the predicted models suggests AlphaFold3 and ColabFold with templates as the most reliable predictors. Additionally, it demonstrates that compared to ColabFold, AlphaFold3 predictions are more consistent in regards to whether a model is correct or incorrect. This is indicated by the significantly higher proportion of cases where all five models in a prediction were classified as incorrect and the lowest proportion where only one model is incorrect.

Moreover, the consensus of prediction quality across the three datasets (CF-T, CF-F and AF3) was analyzed by comparing models of the same rank (Fig. 1C(i), (ii), Suppl. Fig. 2). When the ‘high’, ‘medium’ and ‘acceptable’ quality fractions are combined (considered ‘correct’ based on CAPRI criteria), the models predicted by the three different methods are comparable in terms of quality in 78.4% of cases (Venn diagram in Fig. 1C(i)). The highest proportion (12.9%) of uniquely correct predictions was observed for AF3, suggesting that running AlphaFold3 may enhance the likelihood of generating reliable models. However, Venn diagrams illustrating just high-quality models (based on CAPRI criteria) reveal that only 57.1% of the cases were consistent across all three datasets (Suppl. Fig. 2). Notably, among high-quality models, AF3 achieved the highest proportion of unique predictions (16.8%). This suggests that using AlphaFold3 may increase the likelihood of generating high-quality models. The Venn diagram shows that CF-F had the highest proportion of incorrect predictions (10.4%) based on CAPRI criteria (Fig. 1C(ii)), consistent with its highest proportion of incorrect models in the benchmark set (Fig. 1A). The acceptable fraction showed the lowest agreement (14.0%) between the three prediction methods (Suppl. Fig. 2), suggesting that in cases where predictions are near the threshold for incorrect predictions (DockQ < 0.23), running all three methods and comparing results may yield the most optimal outcome.

To investigate factors contributing to consistent mispredictions and prediction success, we analyzed both the interface size and interface flexibility of targets that were incorrectly predicted across all three datasets, compared to those consistently predicted with high quality. The analysis reveals that interface flexibility, calculated as the average B-factor of the interface, was significantly higher in the mispredicted group (mean for incorrect predictions: 48.30; mean for high quality predictions: 37.55) (Fig. 1D). Additionally, consistently mispredicted targets tend to have smaller interfaces, with the maximum of the distribution shifted toward lower values (incorrect: 32.75; high quality: 60.43) (Fig. 1E), suggesting that larger interfaces may facilitate more reliable predictions. Both differences were statistically significant (Interface size p-value=4.33e-6; b-factor p-value=0.003).

### 3.2. Distributions and correlations of individual prediction-based scores

In order to evaluate the effectiveness of the raw assessment scores (ipLDDT, iPAE, pTM, ipTM, *model confidence*, pDockQ2 and VoroIF, Suppl. Tab. 1) in predicting the quality of the models, these scores were calculated for each model in the three distinct datasets: CF-T, CF-F and AF3. Given the variability in defining the interface based on different distance cutoffs and different types of atoms and positions, the impact of the distance parameter on ipLDDT and iPAE was investigated (Methods). The results indicated no substantial difference in interface detection or ipLDDT and iPAE calculations across the various distance parameters (Suppl. Fig. 3). This suggests that the distance parameter has a marginal influence on these scores.

Consequently, a heavy atom distance cutoff of 5 Å was adopted for the definition of the interface and the calculation of the ipLDDT and the iPAE in agreement with the interface threshold used by AlphaFold to define the interface (Evans et al. 2022).

Subsequently, the empirical distributions of the scores and the correlation between the scores were analyzed (Fig. 2, Suppl. Fig. 4,5). A common trend across these distributions was the predominance of high scores (low scores for iPAE and PAE), suggesting a generally high quality of the prediction sets, consistent with the findings discussed in 3.1. As expected, *model confidence* exhibited the highest correlation with ipTM across all three datasets, as it is the dominant component of the former. Conversely, pDockQ2 demonstrated the lowest correlation with other scores. Overall, the global scores pLDDT and PAE, had a lower correlation with all other scores. In contrast, the interface-specific scores, ipLDDT and iPAE, had a higher correlation with other scores. This could be due to the interface specificity of most of the other scores (except pTM).

**Fig. 2:**
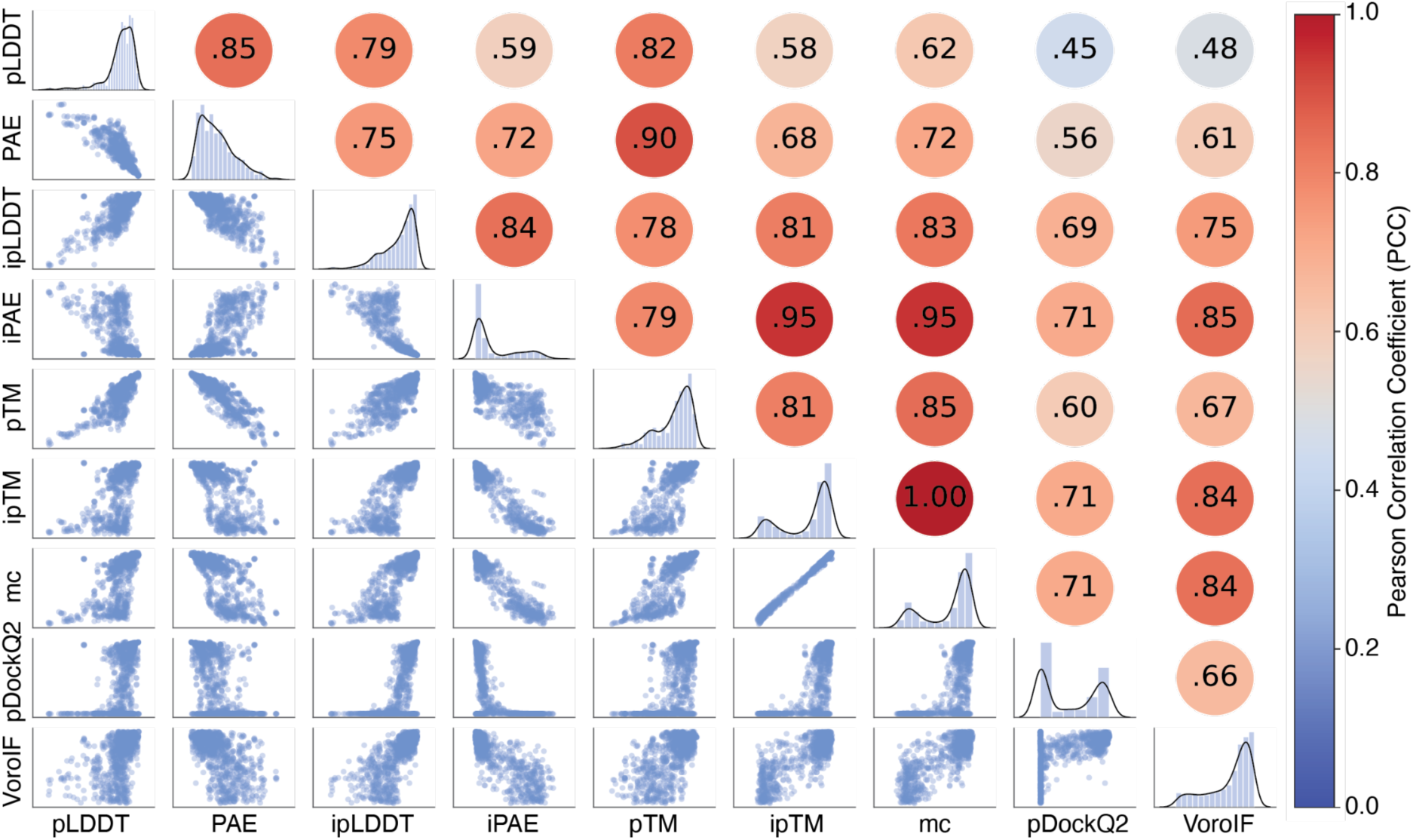
Distribution and correlation of the individual scores (pLDDT, PAE, ipLDDT, iPAE, pTM, ipTM, *model confidence* (mc), pDockQ2 and VoroIF) for the CF-T dataset. The lower-left section displays the pairwise correlation between the scores. The diagonal shows the distribution of each raw score within the benchmark set. The upper-right section provides the PCC between the scores with red indicating high and blue low PCCs.

Overall, the individual prediction-based scores were in good agreement with our observation that these datasets comprised a high proportion of high-quality models. Moreover, the described trends were consistent across all three datasets (CF-T, CF-F and AF3) (Fig. 2, Suppl. Fig. 4,5).

### 3.3. Correlation of individual prediction-based scores with DockQ

To evaluate the effectiveness of individual scores in assessing the quality of the predicted models in the three datasets, PCC between these scores and DockQ were calculated. The results were consistent across CF-T, CF-F, and AF3, with mean PCCs of 0.70, 0.72, and 0.64 (averaged over the individual scores), respectively. However, Wilcoxon signed-rank test showed that only the differences between both ColabFold datasets and AF3 were statistically significant (p-value=0.004). These findings suggest that the scores were more effective in evaluating models generated by ColabFold without templates than by AF3. Notably, despite the generally lower quality of models in the CF-F dataset (Fig 1A) compared to CF-T and AF3, the assessment scores performed best for these models (Fig. 3A). ipTM and *model confidence* showed the highest correlations with DockQ across all datasets (Suppl. Fig. 6), highlighting them as the most reliable single-score metrics for assessing protein complex models, regardless of the prediction method used.

**Fig. 3:**
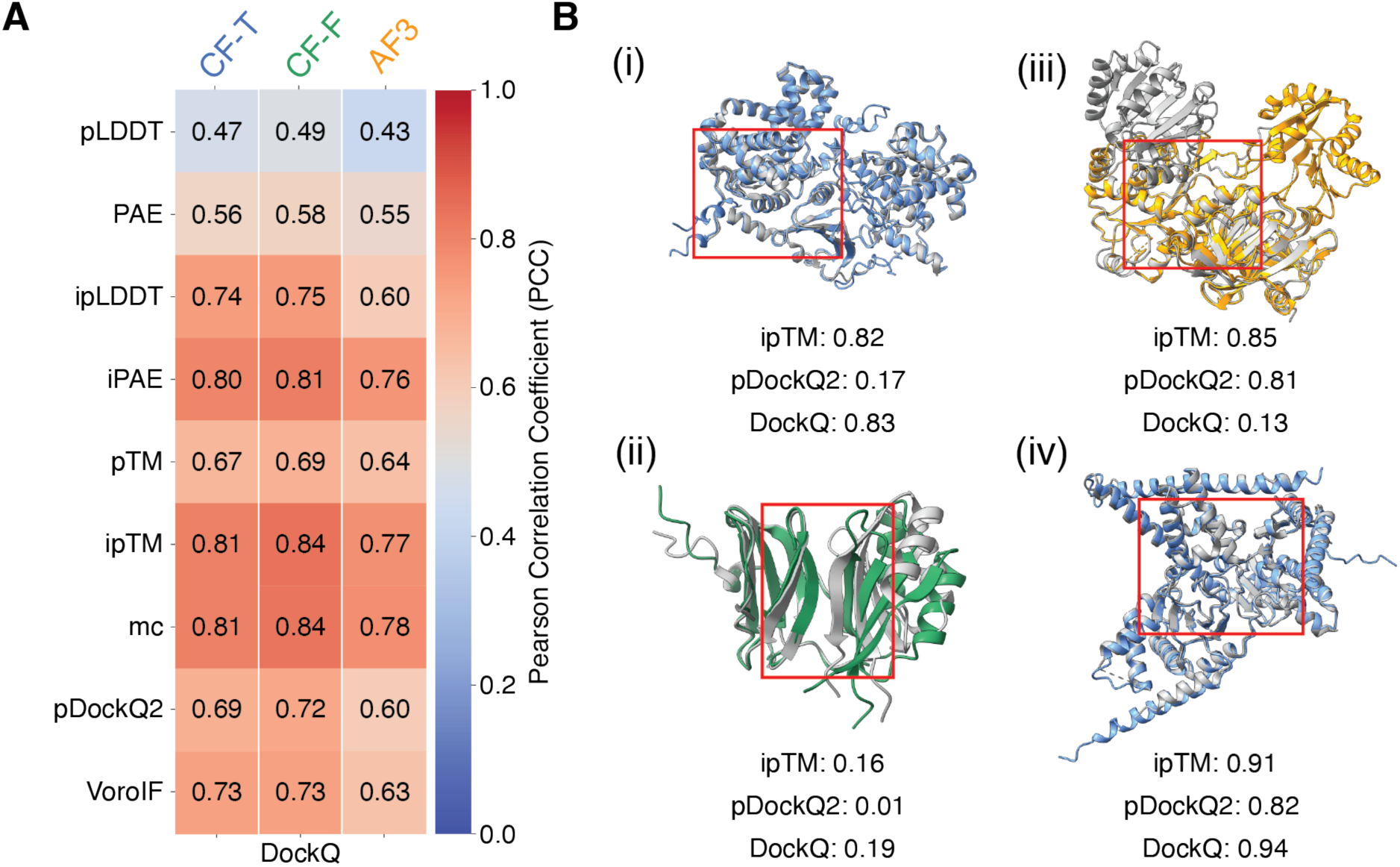
Quality of the individual scores and representative structural predictions with scores. **A:** PCC of the individual scores (pLDDT, PAE, ipLDDT, iPAE, pTM, ipTM, *model confidence*, pDockQ2 and VoroIF) with DockQ. The colorbar is a gradient from blue to red indicating low to high correlation, respectively for CF-T, CF-F and AF3. **B(i):** CF-T prediction (blue) and target structure (PDB: 9H8S) (gray). **B(ii):** CF-F prediction (green) and target structure (PDB: 8B2R) (gray). **B(iii):** AF3 prediction (orange) and target structure (PDB: 7PPZ) (gray). **B(iv):** CF-T prediction (blue) and target structure (PDB: 7PAY) (gray). In **B(i-iv),** the interface region is marked by a red box.

Furthermore, the interface-specific scores, ipLDDT and iPAE, outperformed their global versions, pLDDT and PAE, as they exhibited a significantly higher correlation with the DockQ. Fig. 3B(i) illustrates a case (PDB: 9H8S) where ipTM correctly predicts a high-quality model (DockQ=0.83), whereas pDockQ2 suggests the opposite. In contrast, Fig. 3B(ii) presents an example (PDB: 8B2R) where both ipTM and pDockQ2 correctly reflect low model quality (DockQ=0.19). Fig. 3B(iii) highlights an incorrect assessment by both ipTM and pDockQ2 for a low-quality prediction, due to wrong domain organisation resulting in incorrect interface prediction (DockQ=0.13). Finally, Fig. 3B(iv) shows a case where both metrics correctly agree on the high quality of the model (DockQ=0.94) although ipTM is more confident.

#### Determinants of high- and low-quality models

The individual scores provide only the absolute values without any established thresholds, thus limiting their application for binary classification of the model as correct or incorrect. To address this limitation, we performed Receiver Operating Characteristic (ROC) curve analyses for each score, allowing the determination of cutoffs based on (Melo and Sali 2007) (Methods).

For defining the ground truth, models labelled as ‘incorrect’ by DockQ classification were defined as incorrect models, while those falling into the ‘acceptable’, ‘medium’, and ‘high’ DockQ categories were considered as “correct” predictions. Using ROC curves, optimal thresholds were derived for each dataset (Fig. 4A).

**Fig. 4:**
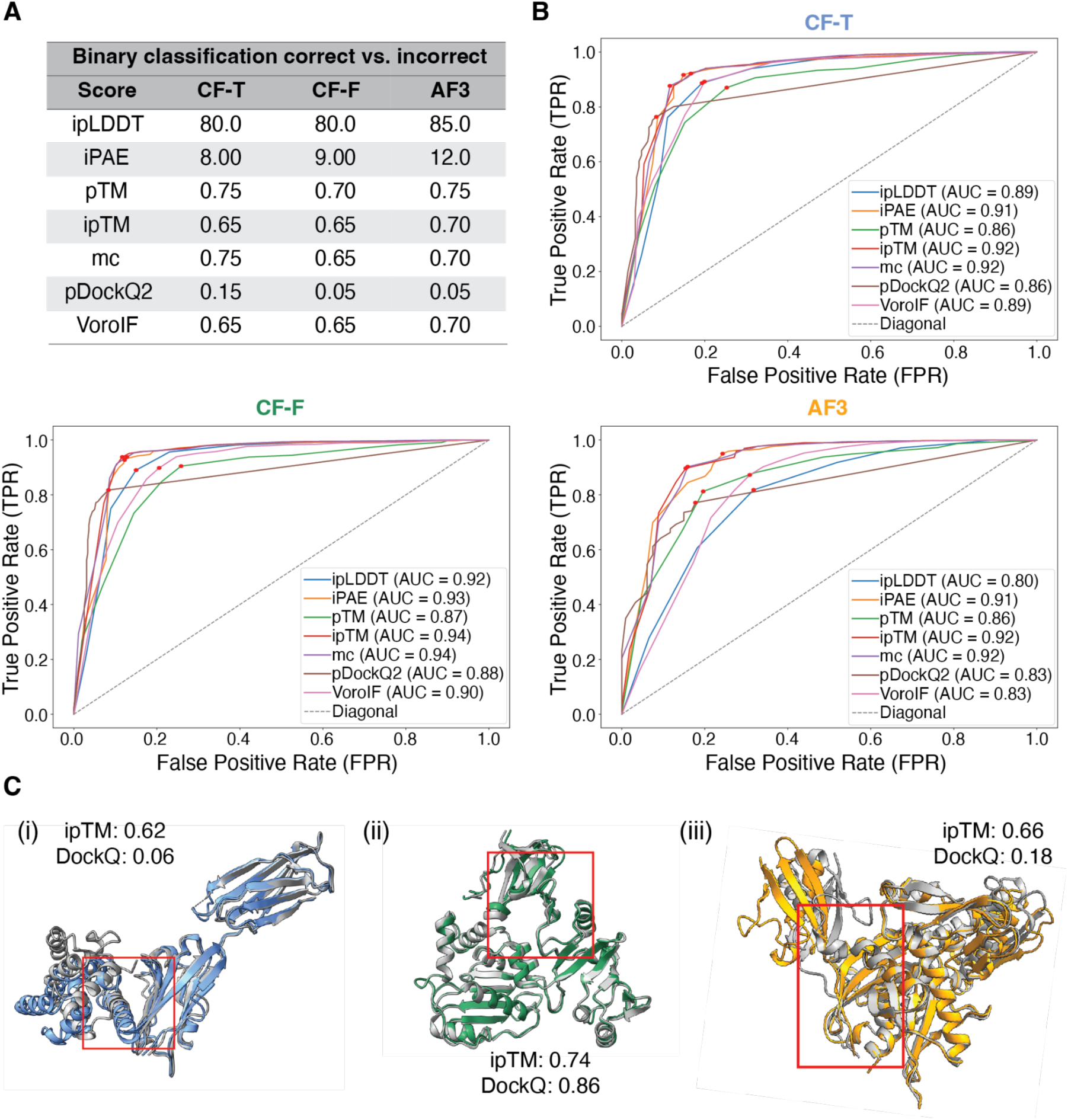
ROC curves, determined cutoffs and example predictions. **A:** Determined thresholds for the scores distinguishing correct (‘high’, ‘medium’, ‘acceptable’ DockQ) and incorrect (‘incorrect’ DockQ) models for CF-T, CF-F and AF3. **B:** ROC-curves and AUC of the raw scores (ipLDDT, iPAE, pTM, ipTM, *model confidenc*e, pDockQ2 and VoroIF) for CF-T, CF-F and AF3. The binary classification is based on the DockQ classification correct (‘high’, ‘medium’, ‘acceptable’ DockQ) and incorrect (‘incorrect’ DockQ) **C(i):** CF-T prediction (blue) and target structure (PDB: 7XYQ) (gray). **C(ii):** CF-F prediction (green) and target structure (PDB: 7EGS) (gray). **C(iii):** AF3 prediction (orange) and target structure (PDB: 7FGM) (gray). In **C(i-iii)** interfaces are marked by a red box.

Additionally, Area Under the Curve (AUC) (Fig. 4B), accuracy and precision (Suppl. Table 2) were calculated to assess the ability of the thresholds to distinguish between models of low and high accuracy as designated by DockQ (Methods). While the precision values of all three datasets were similarly high, the AUC and accuracy values for the AF3 dataset were lower than those for CF-T and CF-F. This result is consistent with Section 3.3, which shows comparatively lower PCC for AF3. Notably, the optimal cutoffs determined for pDockQ2 were unexpectedly low given its scoring range of 0 to 1. On the other hand, the highest AUC (Fig. 4B) and accuracy (Suppl. Table 2) values were observed for the ipTM and *model confidence* scores, in agreement with the findings of Section 3.4.

According to the guidelines in the frequently asked questions of the AlphaFold Server (Accessed October 29, 2024. https://alphafoldserver.com/faq). ipTM > 0.8 indicates high-quality predictions, ipTM < 0.6 suggests likely failed predictions, and ipTM in 0.6 - 0.8 range falls into a “gray zone” where predictions could be correct or incorrect.

However, the importance of providing a specific cutoff to separate correct from incorrect models arises when the model’s ipTM falls within the gray zone. In all three examples shown in Fig. 4C, the ipTM scores fall within the gray zone defined in AlphaFold server, yet the outcomes vary significantly. In Fig. 4C (i and iii), the predictions by CF-T (target PDB: 7XYQ) and AF3 (target PDB: 7FGM) do not pass the ipTM corresponding cutoffs (0.65 and 0.70, respectively) and are categorized as ‘incorrect’ based on DockQ scores.

In contrast, Fig. 4C(ii) shows a CF-F prediction (target PDB: 7EGS for which the ipTM score surpasses our cutoff of 0.65, with the DockQ score classifying this prediction as of ‘high’ quality. Therefore, for the subset of structures of CF-T, CF-F and AF3 with ipTM values in the range of 0.6-0.8, we evaluated the accuracy with cutoffs determined by ROC analyses (CF-T: 0.65; CF-F: 0.65; AF3: 0.70, Fig. 4B). This allowed us to differentiate between correct and incorrect models even in the gray zone with accuracies of 0.82, 0.86, 0.87 for CF-T, CF-F and AF3, respectively (Suppl. Table 3).

### 3.4. Combined scoring and testing

In addition to evaluating the individual scores and their discriminatory power, a combined score was developed to improve the assessment of predicted protein complex models. Various approaches were tested to establish a combined score (Methods). Given that *model confidence* and pDockQ2 are computed by combining other evaluated scores, these scores were excluded.

A linear regression-based model for the combination of all scores (ipLDDT, iPAE, pTM, ipTM, and VoroIF) was consistently identified as the best-performing (based on PCC) combined score across all three datasets (CF-T, CF-F and AF3), and referred to as C2Qscore (Combined Complex Quality score). This score was selected for further analyses (Suppl. Table 4, for weights, bias and maxima see Suppl. Tab. 5-7). Wilcoxon signed-rank test indicated that C2Qscore provided a statistically significant improvement compared to the unweighted combination of the individual scores (Suppl. Table 8).

Finally, ROC curve analysis was conducted to determine a cutoff for differentiating between correct models and incorrect ones (Suppl. Table 9). This analysis yielded a cutoff of 0.55 (AUC: 0.92) for models predicted using ColabFold with templates, 0.48 (AUC: 0.94) for models predicted without templates, and 0.52 (AUC: 0.92) for models predicted by AF3.

To further assess the performance of C2Qscore, it was tested on an independent dataset of 39 heterodimeric protein complexes determined by X-ray crystallography (resolution better than 3 Å), which were randomly selected during the construction of the benchmark dataset and excluded from it to serve as an independent test set. DockQ and all prediction-based scores (including the combined ones) were calculated on 195 models for each of the datasets (CF-T, CF-F, and AF3). The cutoffs defined in Sections 3.5 and 3.6 were applied to all scores to calculate the accuracy, precision, recall and specificity (Fig. 5). C2Qscore showed the same model classification accuracy as the unweighted score (simple average of the individual scores) for CF-T. For CF-F and AF3, C2Qscore achieved slight improvements in all four metrics compared to the unweighted combined scores. When compared to the individual scores, C2Qscore performs equally well or better than nearly all individual scores across CF-T, CF-F, and AF3. Exceptions were observed for iPAE and ipTM in CF-T, where precision and specificity matched those of C2Qscore, but accuracy and recall were slightly higher for these individual scores (Suppl. Fig. 7).

**Fig. 5:**
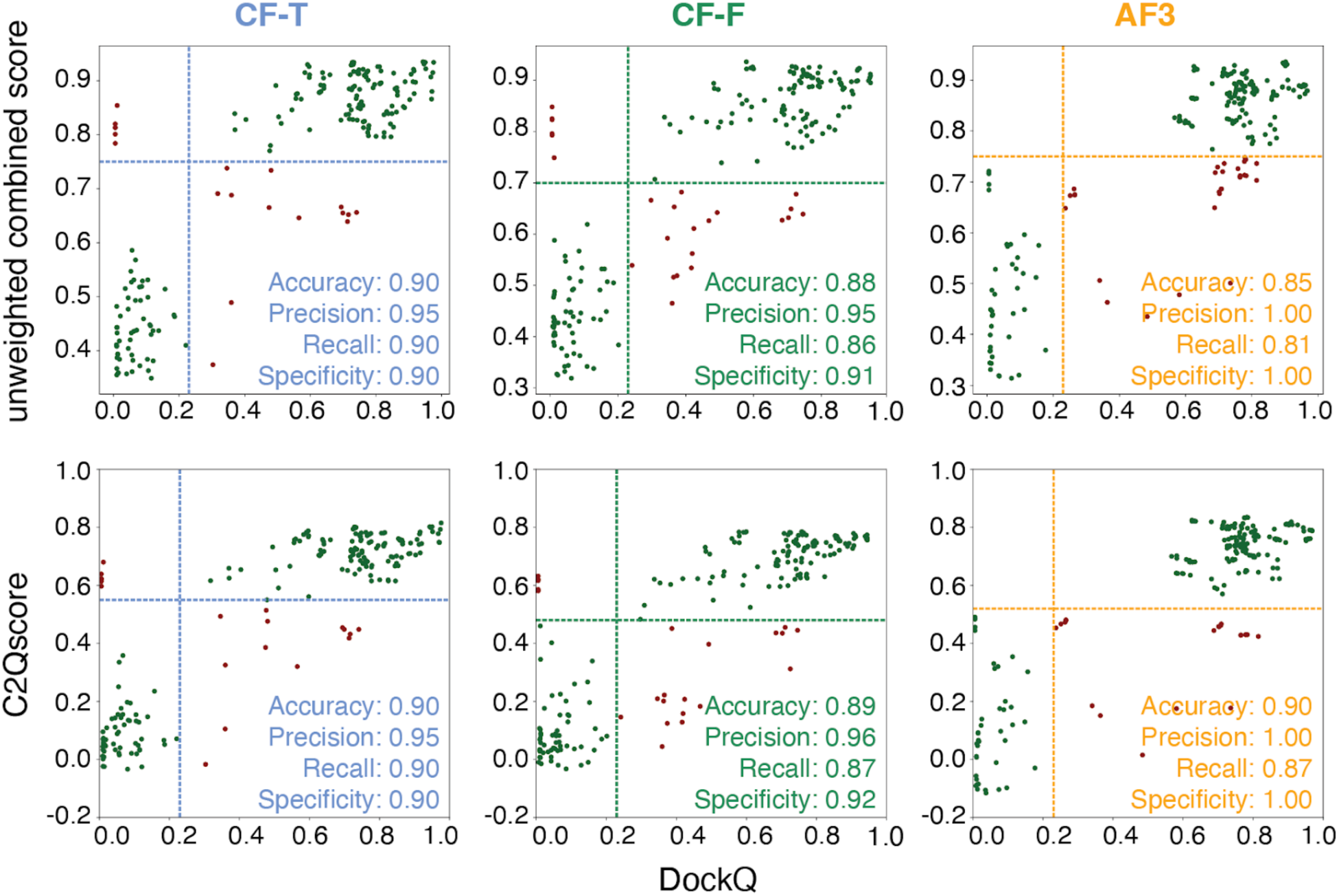
Unweighted combined score and C2Qscore for the X-ray crystallography test set for CF-T, CF-F and AF3. True positives and true negatives are plotted in dark green and false positives and false negatives in dark red. The cutoffs determined in Section 3.6 are shown as dotted lines in the plots. C2Qscore ranges from −0.45 to 0.88 for CF-T, −0.37 to 0.87 for CF-F and −0.33 to 0.93 for AF3.

### 3.7 Evaluating the scores on dimers from larger assemblies

Next, we evaluated the performance of the unweighted combined score and C2Qscore on a dataset of dimers derived from assemblies of higher order (≥ trimeric). This evaluation aimed to assess the performance of the prediction methods and the scoring approaches on dimers, without prior knowledge of the biological context. The cutoffs determined in Section 3.5 were applied. This set included a total of 156 dimeric target structures derived from cryoEM assemblies at resolutions higher than 2.3 Å. DockQ and all prediction-based scores (including the combined ones) were computed for 768 models for CF-T, 769 models for CF-F, and 720 models for AF3.

Here, as before, the performance of C2Qscore was similar to the unweighted score, showing a mild improvement in model classification based on DockQ for CF-T and AF3, while a mild decrease in performance of CF-F (Fig. 6, Supp. Fig. 7). The scoring of these models had a higher percentage of false negatives across all datasets compared to the X-ray test set (Fig. 5, Fig. 6A, Suppl. Fig. 8).

**Fig. 6:**
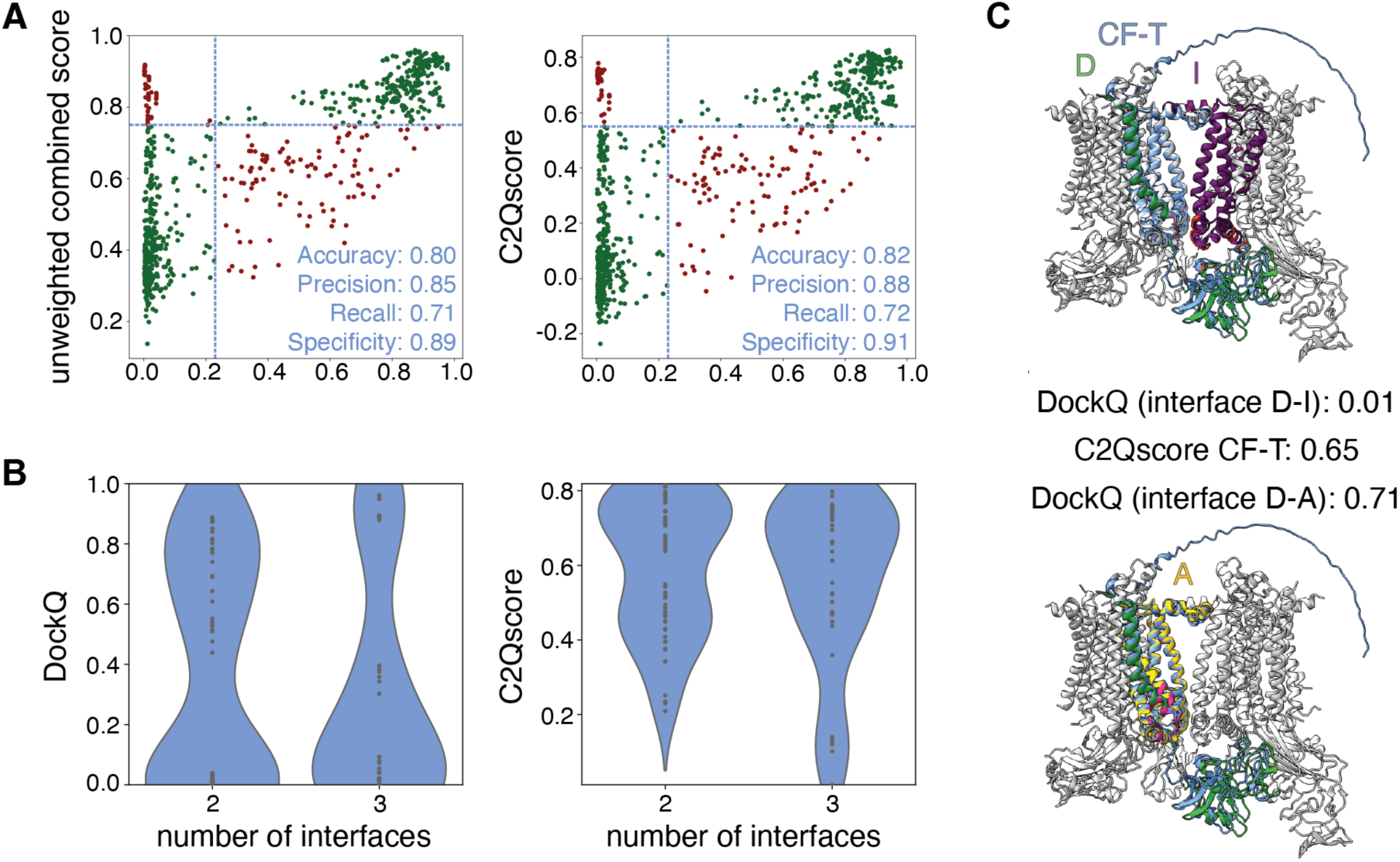
**A:** Unweighted combined score and C2Qscore for the large assembly test set for CF-T. True positives and true negatives are plotted in dark green (upper right and lower left) and false positives and false negatives in dark red (upper left and lower right). The cutoffs we determined based on the benchmark set are shown in light blue. **B:** Distribution of DockQ and C2Qscore for CF-T cases of two (n=58) or three (n=42) potential configurations, for which the prediction resembles one of the configurations. **C:** CF-T prediction superposed on the large assembly (PDB: 9ES7; chains D&I). The prediction (blue), the target chains D (green), I (purple) and the homologue of chain I, chain A (gold), are highlighted on the assembly (gray). The interface of chain D and I (red, top, target) and chain D and A (hotpink, bottom, other possible configurations) are shown. The DockQ for the alignment of the prediction with the complex D-I and D-A and C2Qscore for the prediction are reported.

To understand the general decrease in accuracy compared to the X-ray test set, we further analysed the cryoEM models by investigating the distribution of DockQ and C2Qscore in relation to the number of possible configurations of the dimeric complex derived from large assemblies. In some cases, the target chains of a dimer, which is part of a larger complex, has homologous counterparts within the assembly forming different configurations (Fig. 6B). Since AlphaFold2 and AlphaFold3 only accept the sequence as input, one cannot know in advance which of the available configurations correspond to the predicted structure. Fig. 6C depicts the dimeric complex formed by the chains D and I of the large assembly with PDB-code 9ESY. Chain I is homologous to chain A in the assembly, which also interacts with chain D, resulting in two possible configurations for the target complex. The CF-T prediction closely resembles the dimer formed by chains A and D (DockQ 0.71), but because DockQ is calculated for the complex formed by chains I and D, DockQ is very low (0.01). In contrast, C2Qscore for this prediction is relatively high, at 0.65.

In a typical prediction scenario, where no ground truth is available, the number of potential configurations of a complex is unknown. Therefore, it is crucial to assess the reliability of the scores when various spatial arrangements for two chains to form a dimer (“configuration”) exist. To address this, we used the “large assembly set” and analyzed the DockQ and C2Qscore versus the number of configurations per target dimer for the entire “large assembly” set. The dataset was filtered to include only those templates for which more than one dimeric configuration exist, specifically those with two or three configurations. Furthermore, the predictions were required to be classified as correct based on DockQ (DockQ ≥ 0.23) for one of the existing configurations. It is important to note that the presented DockQ scores (Fig. 6B) correspond to the alignment between the target structure and the prediction. This, by definition, will result in low DockQ values in cases where the prediction closely resembles one of the alternative configurations but is compared to the target configuration.

For cases with two potential configurations of the dimer (n=58) the DockQ showed a bimodal distribution, featuring very high and very low scores (Fig. 6B). However, the C2Qscore distribution differed significantly, with many models scored with very high or medium-to-high scores. For cases involving three possible configurations in CF-T (n=42), the DockQ score peaked at low scores (due to more “different” configuration from the target), while the distribution of the C2Qscore leaned towards higher scores. Similar trends were observed in the CF-F and AF3 dataset (Suppl. Fig. 9).

These findings suggest that both AF2 and AF3 struggle to differentiate between probability and certainty when multiple configurations of a dimer are present. As a result, lower scores (medium-to-high or lower) may be assigned to highly accurate predictions, making the scores less reliable. This could explain the general decrease in accuracy of the models in the large assembly set compared to the X-ray test set.

### 3.8 C2Qscore as a tool

The algorithm for calculating the C2Qscore for AlphaFold2/ColabFold and AlphaFold3 models has been integrated as a tool into our ChimeraX plug-in PICKLUSTER v.2.0 (Pettersen et al. 2021; Genz et al. 2023). The new PICKLUSTER v.2.0 identifies and clusters protein interfaces based on distance, providing a targeted approach for their analysis. It offers access to a range of scoring metrics, including the following scores analyzed in this work: pLDDT, PAE, ipLDDT, iPAE, pTM, ipTM, and *model confidence*. Here, PICKLUSTER v.2.0 has been updated to support predictions generated by AlphaFold3, and the newly developed C2Qscore has been integrated as a feature in the user-interface. To further enable straightforward application of our method, we have also added the calculation of the C2Qscore as a command-line tool, which is available at https://gitlab.com/topf-lab/c2qscore.

## 4. Discussion

In this study, we presented a comprehensive evaluation of commonly used scores for assessing protein complex models and compared three prediction methods: ColabFold with templates (CF-T), ColabFold without templates (CF-F), and AlphaFold3 (AF3). Using a benchmark set of 325 heterodimeric, high-resolution target structures from the PDB, we analyzed the ability of these methods to produce accurate models based on the CAPRI DockQ classifications. We also assessed the effectiveness of the following scores: ipLDDT, iPAE, pTM, ipTM, *model confidence*, pDockQ2 and VoroIF, in distinguishing correct from incorrect predictions.

Overall, we found that ipTM and *model confidence* consistently show the highest PCC with DockQ, highest AUC for classifying models as correct or incorrect and high accuracies across all datasets (CF-T, CF-F, and AF3), suggesting these scores as the most reliable ones to assess heterodimeric protein complex models with both ColabFold and AlphaFold3.

Furthermore, in general the interface-specific scores, ipLDDT and iPAE, showed higher PCCs with DockQ than their global counterparts, pLDDT and PAE, consistent with the findings of (Yin et al. 2022). This highlights the importance of interface-specific metrics for assessing protein complex models.

Interestingly, our results reveal that also iPAE exhibits a high PCC with DockQ, ranking just below ipTM and *model confidence*. Additionally, we observe a strong correlation between PAE and pTM, and between iPAE and ipTM. This can be attributed to the method used to compute ipTM and pTM, both of which are derived from the PAE matrix (Evans et al. 2022). However, during this process, the values within the PAE matrix are weighted, such that residue pairs with lower PAE values contribute more significantly to ipTM and pTM. When averaged across the entire structure, this weighting effect is diminished.

Importantly, the cutoffs we define for differentiating correct from incorrect models using ipTM fall into the gray zone thresholds proposed by the AlphaFold web server (Accessed October 29, 2024, https://alphafoldserver.com/faq). However, while AlphaFold’s ipTM gray zone leaves model quality open to interpretation, our cutoffs offer a practical guide, helping users to assess the reliability of protein complex models based on the chosen structure prediction method and settings. In this study, we establish a new, high-accuracy cutoff specifically for ColabFold, not only for ipTM but for all scoring metrics, to better differentiate between correct and incorrect models.

Notably, the determination of cutoffs was also effective for AF3, though with lower accuracy compared to CF-T and CF-F. Along with lower AUC, accuracy values and lowest mean PCC between the individual scores and DockQ, these findings suggest an overall weaker performance of assessment scores for AlphaFold3 compared ColabFold. This may be due to the differences in the network architecture and confidence estimation of AlphaFold3 compared to AlphaFold2-multimer (Abramson et al. 2024).

According to our benchmark, CF-T and AF3 produce similarly reliable predictions. This contrasts with the findings of (Abramson et al. 2024), which reported that AlphaFold3 improved protein complex accuracy compared to AlphaFold-multimer (v.2.3), the version employed in ColabFold v.1.5.2 (which is also the version used in this work (CF-T)).

Not surprisingly, our results show that as the prediction quality (DockQ) decreases, the agreement among the prediction methods also declines. Interestingly, this suggests that for cases of lower prediction quality, in particular when the scores are of lower prediction quality, using multiple prediction methods as well as evaluation scores and comparison of the results may be beneficial.

We analysed our benchmark for cases where the asymmetric unit (AU) differed from the biological assembly (BA) in the PDB. Interestingly, in such instances, we observed that all three methods could predict either the AU or the BA with high confidence, but without prior knowledge, it would not be possible to determine which outcome reflects the correct structure. This scenario becomes particularly problematic when the predicted AU is correct based on scoring metrics but does not correspond to the functional BA, potentially leading to false interpretations in subsequent experiments. Such cases underline the importance of cautious interpretation of predictions, particularly in scenarios where the biological relevance of the AU and BA is unclear (Schweke et al. 2023).

To improve the ability of assessing protein complex models, we developed a combined score, C2Qscore (Combined Complex Quality score) based on a linear regression model that included ipLDDT, iPAE, pTM, ipTM and VoroIF. C2Qscore enhanced model quality assessment, particularly for CF-F and AF3, and performed comparably for CF-T regarding the unweighted combined scores for the X-ray test set. Its strength lies in integrating multiple scores, which helps reduce reliance on any single metric and improves robustness, especially in ambiguous cases. This is consistent with previous findings showing that combined scoring approaches, such as DockQ itself (Basu and Wallner 2016), consensus-based methods in CASP <u>(Kryshtafovych et al. 2014)</u>, and for assessing protein–protein docking poses <u>(Moal et al. 2013)</u>, tend to offer more robust and generalizable assessments than individual metrics. While individual scores like ipTM or iPAE may perform well in isolation and sometimes outperform C2Qscore, the latter offers greater consistency, particularly in terms of precision and specificity. Given the relatively small size of the test set, these findings should be interpreted cautiously, but they suggest that the combined scoring approach provides a more reliable model classification across different prediction methods.

Testing of C2Qscore on the ‘large assembly’ dataset revealed an increased percentage of false negatives and a reduced classification accuracy across all prediction methods compared to the X-ray test set. This suggests that when multiple configurations of a dimer are possible, AlphaFold2 and AlphaFold3 may give lower scores (medium-to-high rather than high) to highly accurate models, compromising the reliability of the scores.

Finally, it is important to note that this study focused exclusively on protein pairs that are known to interact, using the scoring metrics to evaluate the quality of the predicted complexes. Our goal was to assess how these scoring methods perform in realistic use cases, where predicted complexes often have homologous structures in available databases. Since AlphaFold inherently leverages homology during prediction, excluding such homologs would create an artificial evaluation setting that does not reflect typical user scenarios. Other studies have applied these scores to determine whether a protein pair interacts or not (Schweke et al. 2023). The latter approach should be used with caution, as studies have shown that ipTM is sensitive to variations in sequence constructs, leading to different ipTM scores despite unchanged predicted interface contacts (Bret et al. 2024; Pándy-Szekeres et al. 2024). This suggests that while a high ipTM score indicates confidence in the prediction, a low score does not necessarily imply that protein pairs do not interact and scores in the grey zone may or may not imply interaction. Tools like PPIscreenML (Mischley et al. 2024) may provide potential solutions to address this challenge.

## 5. Conclusion

Our study shows both the strengths and limitations of individual commonly used assessment scores for evaluating heterodimeric protein complex models produced by ColabFold with and without templates and by AlphaFold3. Our PICKLUSTER v.2.0 plug-in and command-line tool enable the usage of these scores along with our combined score, C2Qscore, in an interactive and easy manner. Overall, we recommend ipTM and *model confidence* for general use due to their high correlation with DockQ. However, the interpretation of the scores needs to be performed carefully, particularly for AlphaFold3 models. Notably, our analysis was conducted exclusively on heterodimeric protein complex models without the consideration of multiple interfaces. Further research should investigate the transferability of the results to higher-order complexes. Moreover, exploring different interfaces could reveal how scores perform in situations where interface regions differ significantly in quality. With AlphaFold3 also enabling the prediction of protein-ligand, protein-lipid, and protein-nucleotide interactions, future research could focus on scoring metrics for these types of complexes. This also includes complexes involving antibodies, which were excluded from the analysis in this work. Refining scoring methods and expanding their applicability to diverse complex types and interface characteristics will be for computational protein complex modeling to reach its full potential.

## 6. Methods

### 6.1. Generation of the benchmark dataset (Suppl. Fig. 1)

PDB entries were initially filtered to include hetero-dimeric complexes with a release date after 30.09.2021 (last date used in the AlphaFold training set), a release data before 11.07.2025, and a refinement resolution ≤3.0 Å. The entries were grouped by a sequence identity of >30% using PDB default settings (MMseqs2 (Steinegger and Söding 2017)). The asymmetric unit of representative structures from groups that satisfied the specified criteria were downloaded from the PDB, yielding a total of 671 complexes.

In the subsequent step, interacting complexes were defined as two chains having at least three residues from each chain interacting with residues of the other chain, with heavy atom distance cutoff of 5 Å, using PICKLUSTER (Genz et al. 2023). Non-interacting complexes, as well as any chain with less than 50 residues, were filtered out using the PICKLUSTER pipeline. This process resulted in a total of 376 protein complexes.

In the next step, the antibody database SAbDab (Dunbar et al. 2014) was used to filter out entries containing antibodies or nanobodies. Subsequently, to ensure that the dimers could be classified as heterodimeric complexes, homology within the complexes was assessed by application of CD-HIT (Fu et al. 2012) with a 30% sequence identity threshold, resulting in 270 complexes. Moreover, only templates for which one BA assigned in the PDB equaled the AU were included in the benchmark set, resulting in 223 target complex structures in the final benchmark set.

#### X-ray crystallography test set

During the construction of the benchmark dataset, 39 heterodimeric target structures were randomly selected and excluded from the benchmark set. These 39 structures were used as an independent test set (X-ray test set) for evaluating model performance.

#### Large assembly test set

The set of dimers from larger assemblies (“large assembly test set”) was curated following the protocol established for the benchmark set (Suppl. Fig. 1), with the following modifications to the selection criteria: the dimers were derived from higher-order assemblies (≥ trimeric) with a resolution ≤2.3 Å resolved by cryoEM and a release date after 01.09.2023. A stricter resolution cutoff was implemented compared to the original benchmark set due to the potential variability in local resolution relative to the reported global resolution in structures resolved by cryoEM (Vilas et al. 2020). To ensure the non-redundancy of dimers derived from the same larger assembly, the homology of the dimers was evaluated using CD-HIT (Fu et al. 2012) with a 30% sequence identity threshold, resulting in 156 dimeric target structures included in the large assembly test set.

### 6.2. Model generation

Three AI-based structure prediction approaches were used: CF-F, CF-T and AF3, to create three corresponding datasets.

For the CF-F dataset structural models were predicted from sequence using local ColabFold v1.5.2 (Mirdita et al. 2022). Default parameters were applied for multiple sequence alignment (MSA) generation (MSA mode: mmseqs_uniref_env) and three recycles were performed. For each template, five models were generated, all of which were included in the dataset. Relaxation was performed to all five models.

For the CF-T dataset structural models were predicted using the same criteria as for the CF-F dataset. The only difference was the activation of the template option and the template cutoff date was set to 30.09.2021 to exclude the possibility of finding target PDBs in the templates.

For the AF3 dataset structural models were predicted using the AlphaFold3 web server (https://alphafoldserver.com/) (Abramson et al. 2024) with default settings, which includes the usage of templates with release dates before 30.09.2021.

For all datasets, input sequences used for structure prediction were derived from the corresponding sequence records.

### 6.3. Reference-based score

DockQ v2.0 (Basu and Wallner 2016; Mirabello and Wallner 2024) (https://github.com/bjornwallner/DockQ) (https://github.com/bjornwallner/DockQ) was used to evaluate the quality of the predicted models, by comparing the predicted models of a complex against the native complex structure. The score combines the criteria from the CAPRI competition (Lensink and Wodak 2013) into a single value. Scores range from 0 to 1, with 1 indicating a perfect match between the interface of the predicted model and that of the native complex structure. Threshold values of 0.23, 0.49, and 0.8 were used to categorize predictions as acceptable, medium, and high quality respectively, according to CAPRI standards. DockQ scores were determined by comparing the predicted models of a complex against the native complex structure. DockQ v2.0 was installed according to https://github.com/bjornwallner/DockQ.

### 6.4. Prediction-based scores

The scores used for assessment of models are: pLDDT (predicted local difference distance test) (Jumper et al. 2021), PAE (predicted alignment error) (Evans et al. 2022), ipLDDT (interface pLDDT) (Jumper et al. 2021; Genz et al. 2023), iPAE (interface PAE) (Evans et al. 2022; Genz et al. 2023), pTM (predicted TM-score) (Jumper et al. 2021), ipTM (interface pTM) (Evans et al. 2022), *model confidence* (Evans et al. 2022), pDockQ2 (Zhu et al. 2023) and VoroIF (VoroIF-GNN) (Olechnovič and Venclovas 2023 Jul 21). The details about each score, including the range, what it measures and whether it is interface specific or not, are provided in Supp. Table. 1.

### 6.5. Performance metrics

To measure the overall performance of scores over the different datasets we used a number of approaches:

#### Pearson correlation coefficient

The Pearson correlation coefficient (PCC) gives an indication of the strength of the linear relationship between two datasets. The PCC ranges from −1 to +1 with 0 indicating no correlation and correlations of −1/+1 implying an exact linear relationship. The PCC only detects linear dependencies between the datasets. Positive values indicate a correlation whilst negative values imply anti-correlationPositive values indicate a correlation whilst negative values imply anti-correlation (Benesty et al. 2009).

The PCC was calculated using the stats package provided in the *SciPy* v1.11.3 Python package (Virtanen et al. 2020).

#### ROC-curves and AUC

To identify the best performing cutoff for the scores, a Receiver Operator Curve (ROC) analysis was conducted. The optimal cutoff for the score was determined by maximizing the separation between correctly and incorrectly assessed models, calculated through the True Positive Rate (TPR) and False Positive Rate (FPR), which were plotted as the ROC curve. The best performing cutoff was identified as the point with the maximum distance from the diagonal in the ROC curve (Melo and Sali 2007). For each point on the ROC curve, the cutoff values were incrementally increased depending on the score type (pLDDT/ipLDDT: 5; iPAE/PAE: 1; all other scores: 0.05). The Area Under the Curve (AUC) was calculated to quantify the accuracy of the metric being evaluated, where an AUC of 0.5 indicates random performance, and an AUC of 1 represents perfect discrimination. Both the ROC curves and AUCs were computed using a Python script utilizing the metrics module from the *Scikit-learn* v1.3.0 Python package (Pedregosa et al. 2011).

#### Accuracy

.Accuracy is defined as the ratio of (TP+TN)/(TP+TN+FP+FN), where TP represents the number of true positives, TN the number of true negatives, FP the number of false positives and FN denotes the number of false negatives. The metric is used to evaluate the performance of a model, indicating how frequently the model makes correct predictions. It is determined by comparing the number of correct predictions to the total number of predictions made by the model. The metric is expressed as a value between 0 and 1, with 1 indicating the optimal outcome. The accuracy provides an overall measure of the reliability of a model (Powers 2020).

#### Precision

Precision is defined as the ratio TP/(TP+FP). Precision quantifies the proportion of correctly identified positive samples out of all samples predicted as positive. It therefore reflects the ability of the classifier to avoid falsely classifying negative samples as positive. A precision value of 1 indicates perfect performance, with no false positives, while a value of 0 represents the worst outcome, with all predicted positives being incorrect. Precision shows how often a model is correct when predicting the target class (Powers 2020).

#### Wilcoxon signed rank (WSR) test

The WSR test is a nonparametric test procedure for the evaluation of the significance of paired data (Woolson 2005). This non-parametric test was chosen due to its ability to compare matched data without assuming normality. The p-value was calculated to determine the significance of the observed difference. A significance threshold of p<0.05 was applied. The WSR test was implemented using the stats package provided in the *SciPy* Python package (Virtanen et al. 2020).

### 6.6. Influence of the distance threshold for the definition of the interface on the scores

To determine the interface PAE (iPAE) and interface pLDDT (ipLDDT), the interface was defined as residues with less than 5Å (Evans et al. 2022) distance between heavy atoms (non-hydrogen atoms) of the two chains in the heterodimeric structure, which was calculated with PICKLUSTER (Genz et al. 2023).

To investigate the impact of the distance parameter on ipLDDT and iPAE, we chose to do this analysis on the dataset generated using CF-T, which had 1,612 models in total. We then computed ipLDDT and iPAE on these models using distance thresholds in a range of 5 to 9Å (1Å steps).

### 6.7. Linear and Lasso regression

Linear regression was tested as the first approach for a weighted combined score. It fits a linear model with coefficients by minimizing the residual sum of squares between the target values in the datasets and the ones predicted by the approximation of the linear model. A Lasso regression model was evaluated as the second approach to define a weighted combined score. This approach uses a linear model that is fitted under the assumption that only few of the coefficients are non-zero. Therefore, it identifies the most relevant features and reduces the number of input variables.

Linear and Lasso regression were performed by using the linear_model package provided in the *Scikit-learn* Python package (Pedregosa et al. 2011).

### 6.8. Combined scoring

To enable a combined score, both ipLDDT and iPAE were normalized between 0 and 1, representing the optimal outcome. Subsequently, various approaches were tested to establish a combined score demonstrating superior model assessment capacity. Given that *model confidence* and pDockQ2 are computed by combining other evaluated scores, these scores were excluded.

In the first approach we selected the best combination of scores (ipLDDT, iPAE, pTM, ipTM, and VoroIF) based on PCC with pDockQ as the evaluation criterion. All possible combinations without repetition of equally weighted scores within each set were tested, and PCCs were computed for each. In the second approach we employed linear regression, applied to both the full combination of scores and the best-performing combination identified in the first approach. Finally, Lasso regression was tested as an approach for improvement over using individual scores.

## 7. Acknowledgments and Funding

We thank the Topf group, especially Thomas Mulvaney, and Karen Manalastas-Cantos, for discussions and valuable feedback. This work was supported by the cooperation of Leibniz Institute of Virology Strategic Incentive Program and the Leibniz ScienceCampus InterACt (funded by the BWFGB Hamburg and the Leibniz Association), the BMBF Computational Life Sciences project (ASPIRE, 031L028), the Landesforschungsförderung Hamburg (HamburgX) and DFG CRC1648.

