## Supplementary material for "Assessing scoring metrics for AlphaFold2 and AlphaFold3 protein complex predictions": Suppl.

This supplementary material includes:

- Supplemental text (Page 44)
  - Reasons for calculation failures for scores
- Supplemental figures (Pages 45-52)
  - Suppl. Fig. 1
  - Suppl. Fig. 2
  - Suppl. Fig. 3
  - Suppl. Fig. 4
  - Suppl. Fig. 5
  - Suppl. Fig. 6
  - Suppl. Fig. 7
  - Suppl. Fig. 8
  - Suppl. Fig. 9
- Supplemental tables 1-9 (Pages 53-62)
  - Suppl. Table 1
  - Suppl. Table 2
  - Suppl. Table 3
  - Suppl. Table 4
  - Suppl. Table 5
  - Suppl. Table 6
  - Suppl. Table 7
  - Suppl. Table 8
  - Suppl. Table 9

**Supplemental text**

**Reasons for calculation failures for scores**

The calculation of scores encountered several challenges that led to removal of some of the predicted models from the large assembly test set (Methods). A common issue was the segmentation fault that caused the core being dumped and resulted in unexpected termination. Another issue was the presence of invalid coordinate lines in the .pdb files of templates obtained from the PDB, which resulted in errors when attempting to load the structures in order to calculate the scoring metrics. Moreover, the calculation of scores failed, when PICKLUSTER was not able to detect an interface in the predicted dimeric model (according to the criteria established by PICKLUSTER).

**Supplemental figures**

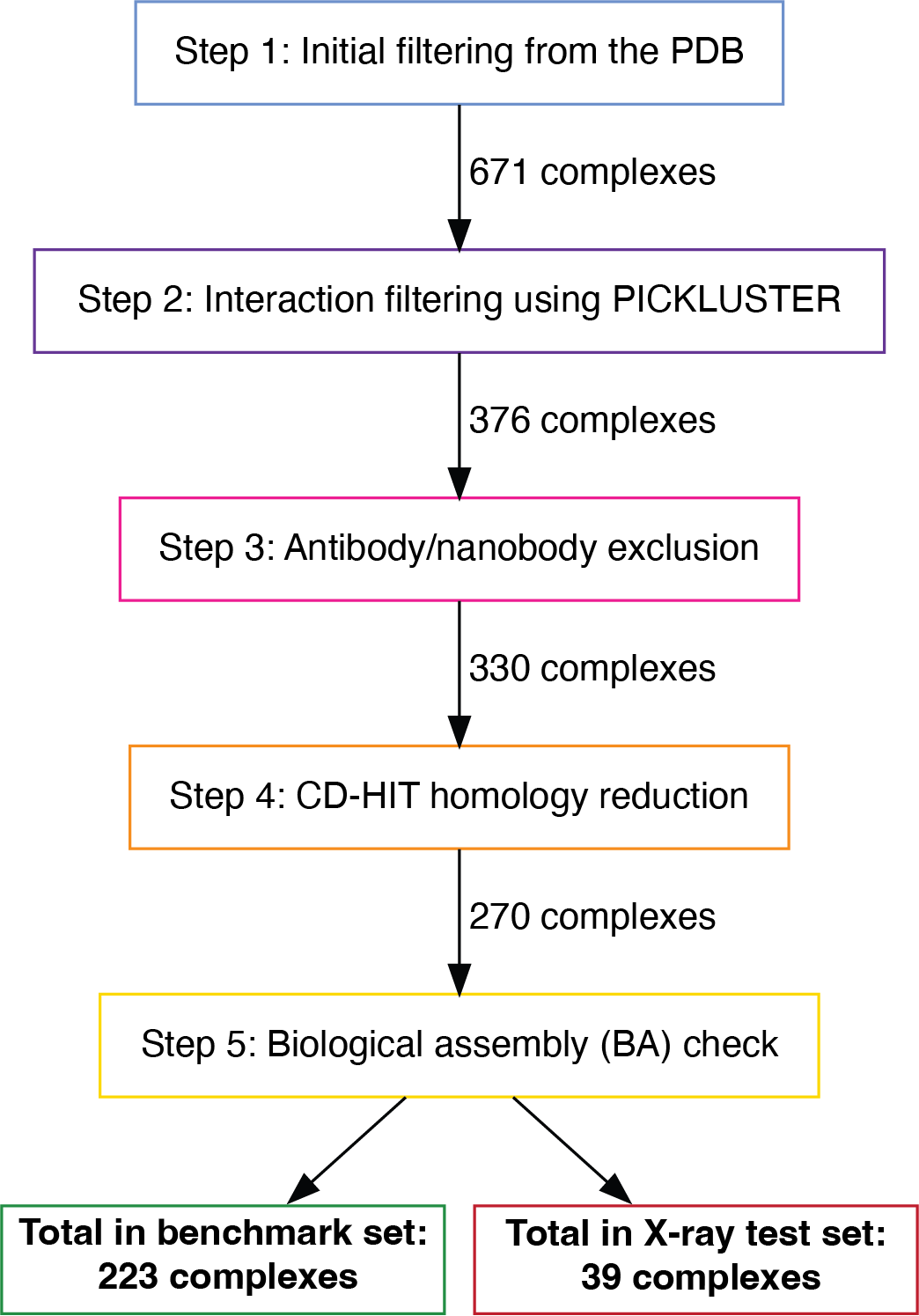

**Suppl. Fig. 1:** Curation pipeline for the benchmark and the X-ray test set.

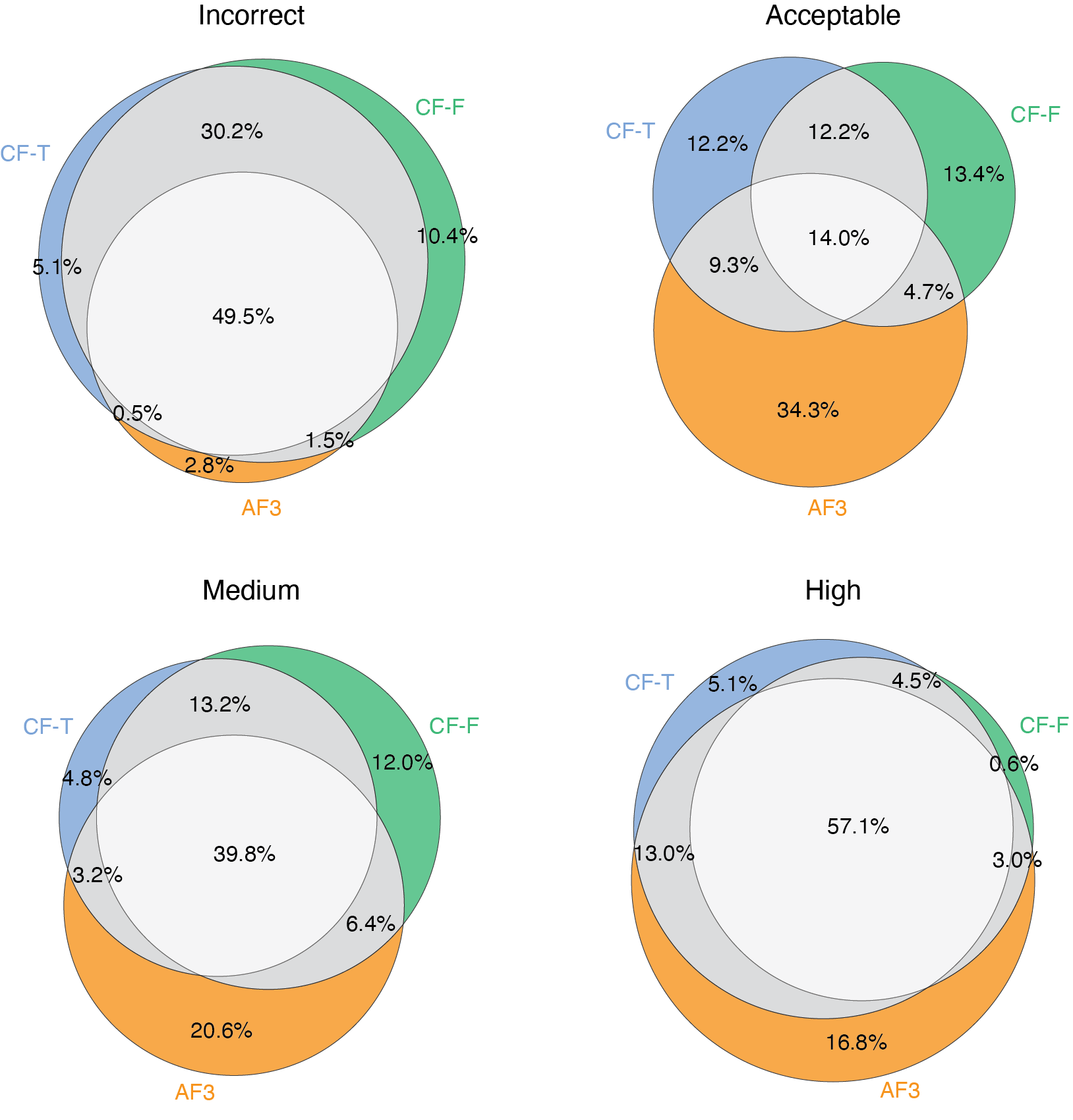

**Suppl. Fig. 2:** Venn-diagrams of CF-T, CF-F and AF3 based on DockQ classifications.

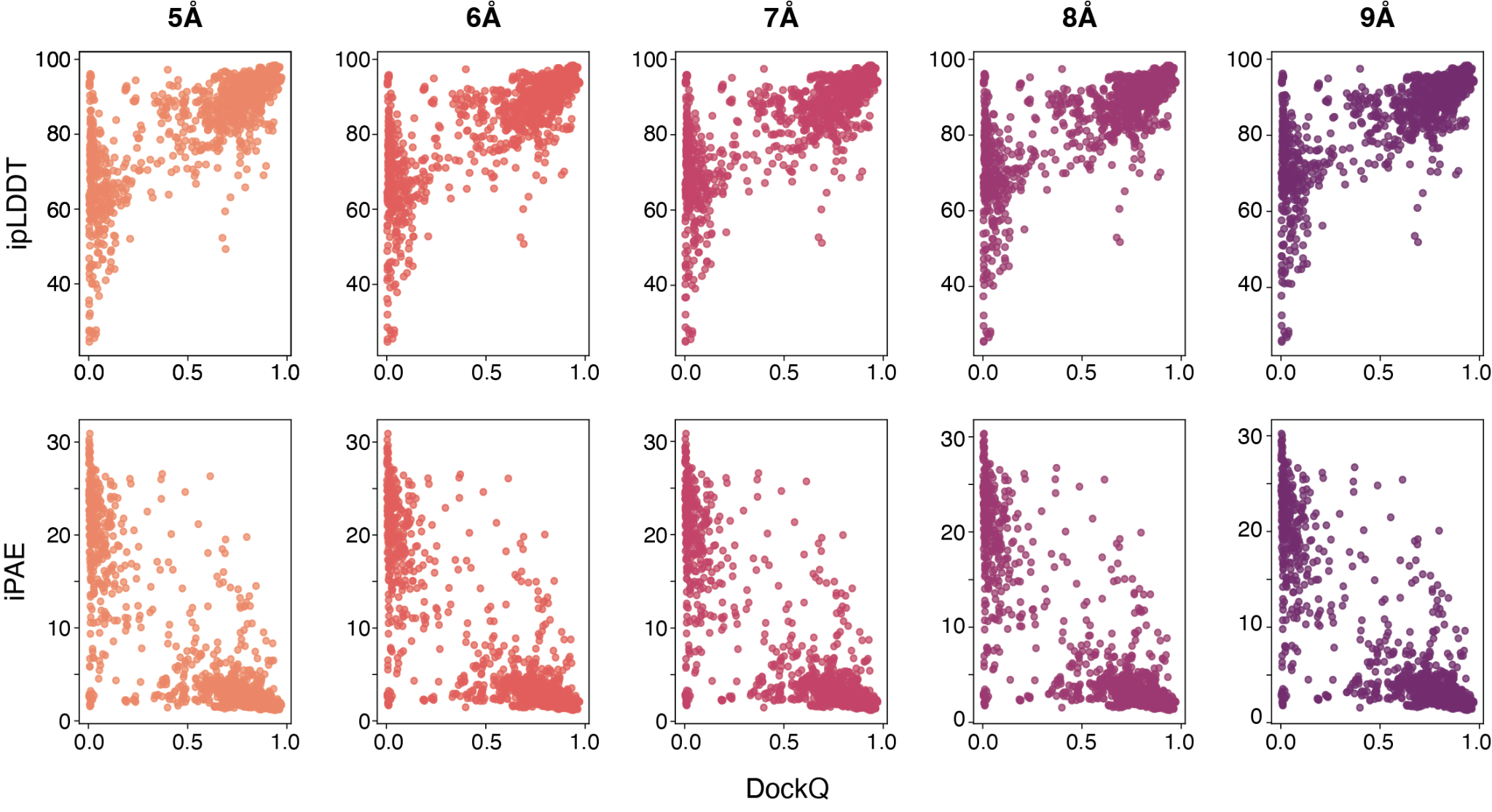

**Suppl. Fig. 3:** Scatter-Plots of the ipLDDT/iPAE vs. DockQ using distance thresholds in a range of 5 to 9Å.

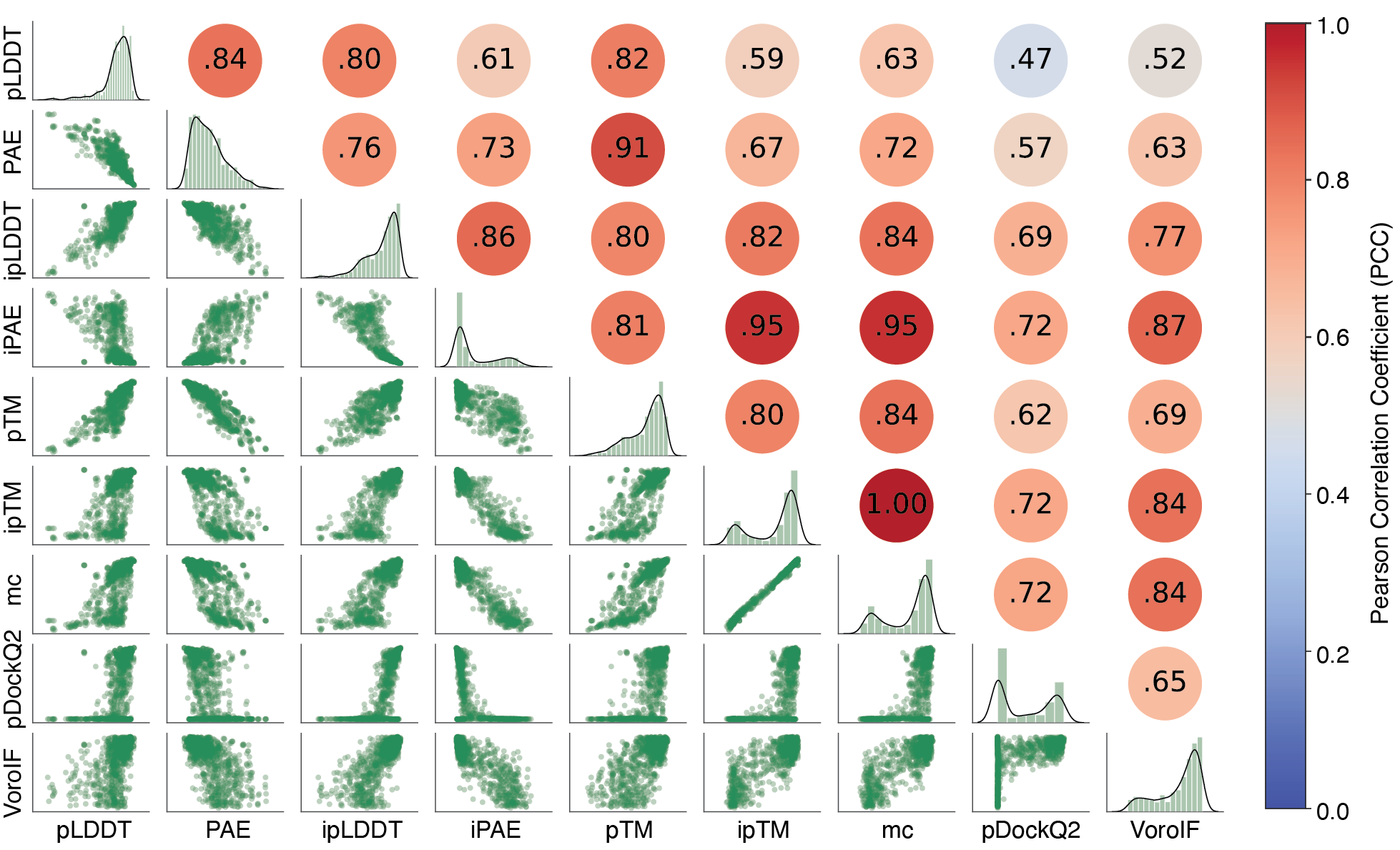

**Suppl. Fig. 4:** Distribution and correlation of the raw scores (pLDDT, PAE, ipLDDT, iPAE, pTM, ipTM, *model confidence* (mc), pDockQ2 and VoroIF) for the CF-F dataset. The lower-left section displays the pairwise correlation between the scores. The diagonal shows the distribution of each raw score within the benchmark set. The upper-right section provides the Pearson correlation coefficients between the scores.

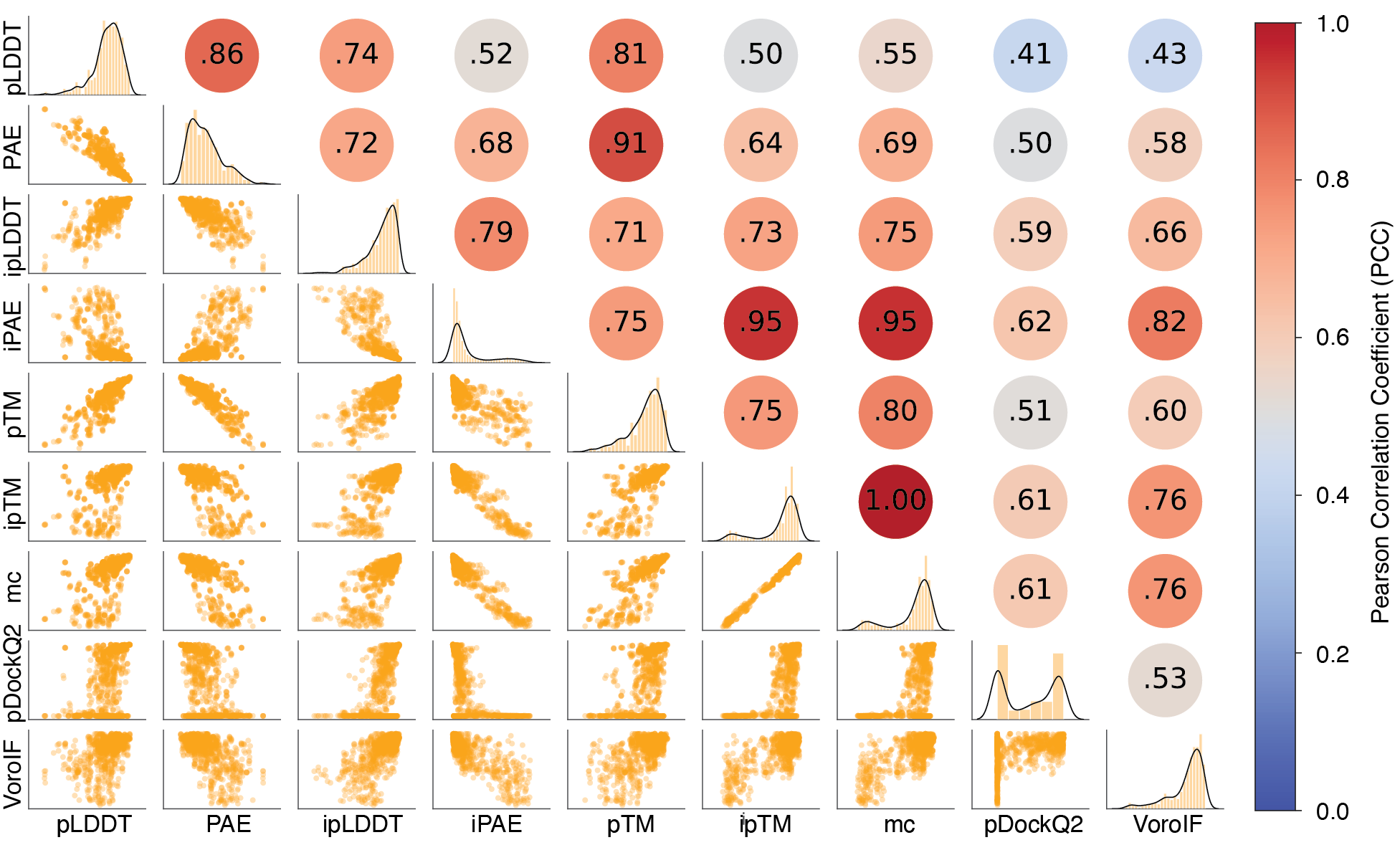

**Suppl. Fig. 5:** Distribution and correlation of the raw scores (pLDDT, PAE, ipLDDT, iPAE, pTM, ipTM, *model confidence* (mc), pDockQ2 and VoroIF) for the AF3 dataset. The lower-left section displays the pairwise correlation between the scores. The diagonal shows the distribution of each raw score within the benchmark set. The upper-right section provides the Pearson correlation coefficients between the scores.

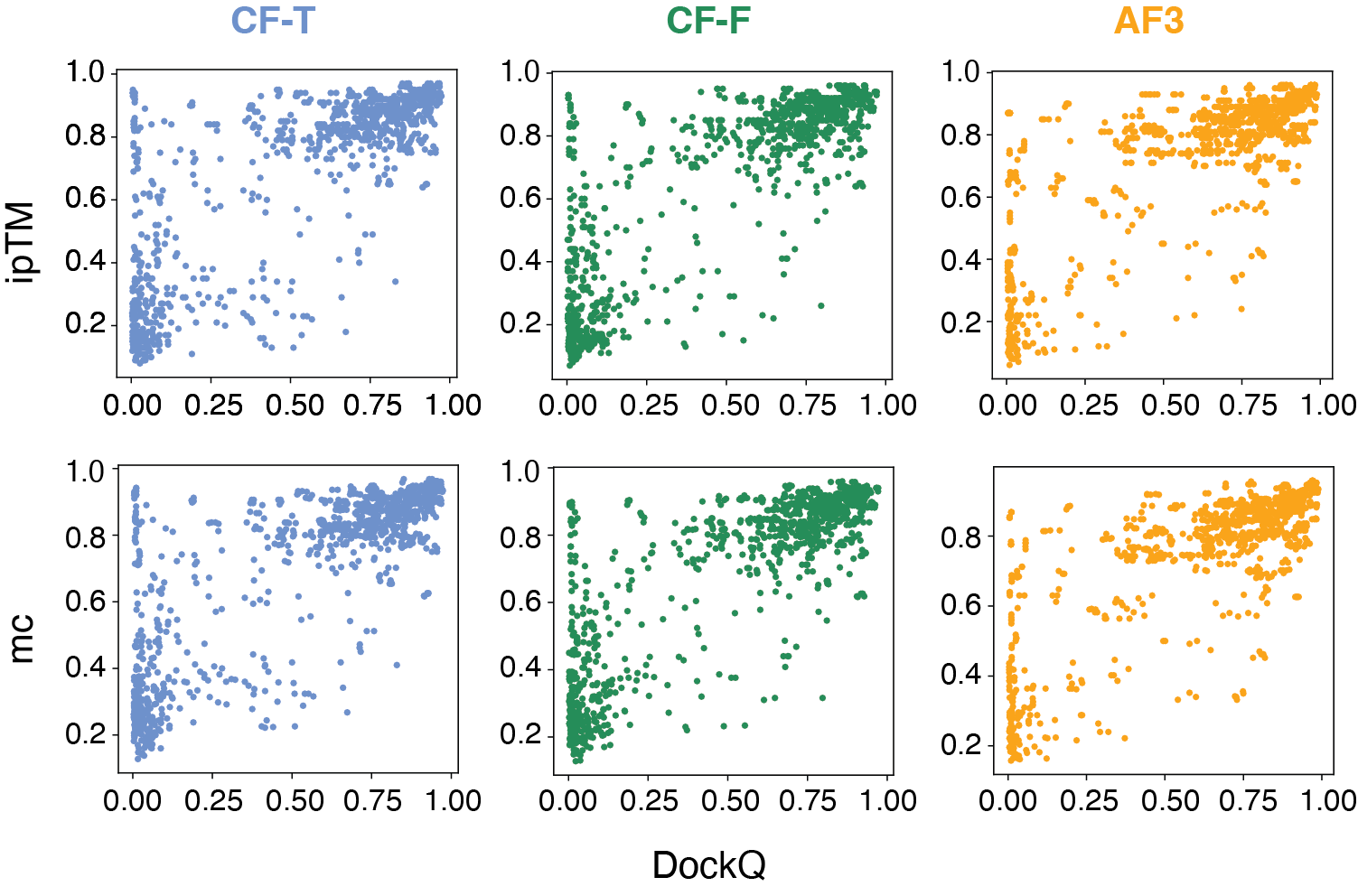

**Suppl. Fig. 6:** ipTM and *model confidence* (mc) vs. DockQ or the benchmark set for CF-T, CF-F and AF3.

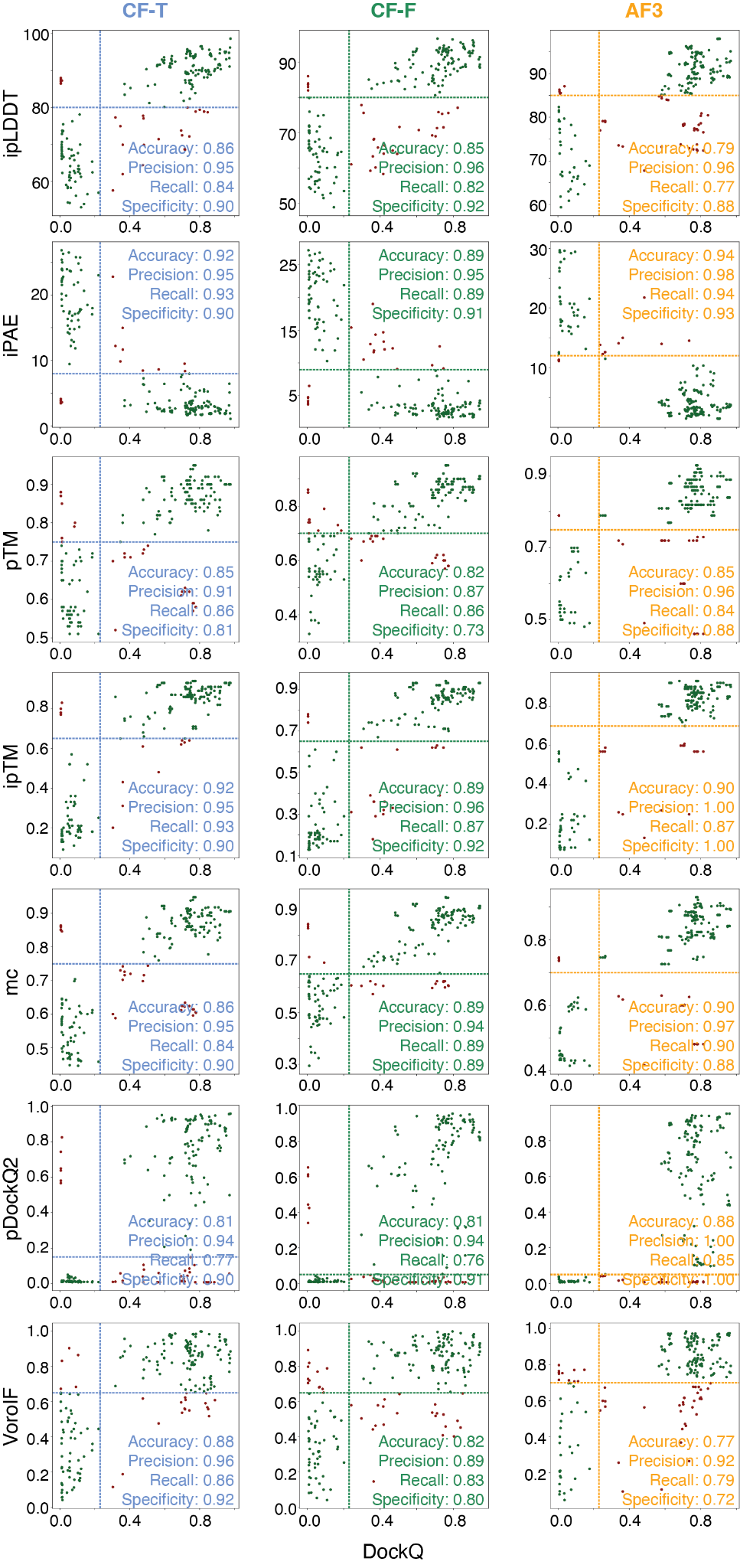

**Suppl. Fig. 7:** ipLDDT, iPAE, pTM, ipTM, *model confidence* (mc), pDockQ2 and VoroIF vs. DockQ for the test set for CF-T, CF-F and AF3. True positives and true negatives are plotted in dark green and false positives and false negatives in dark red. The cutoffs determined in Section 3.5 are shown as dotted lines in the plots.

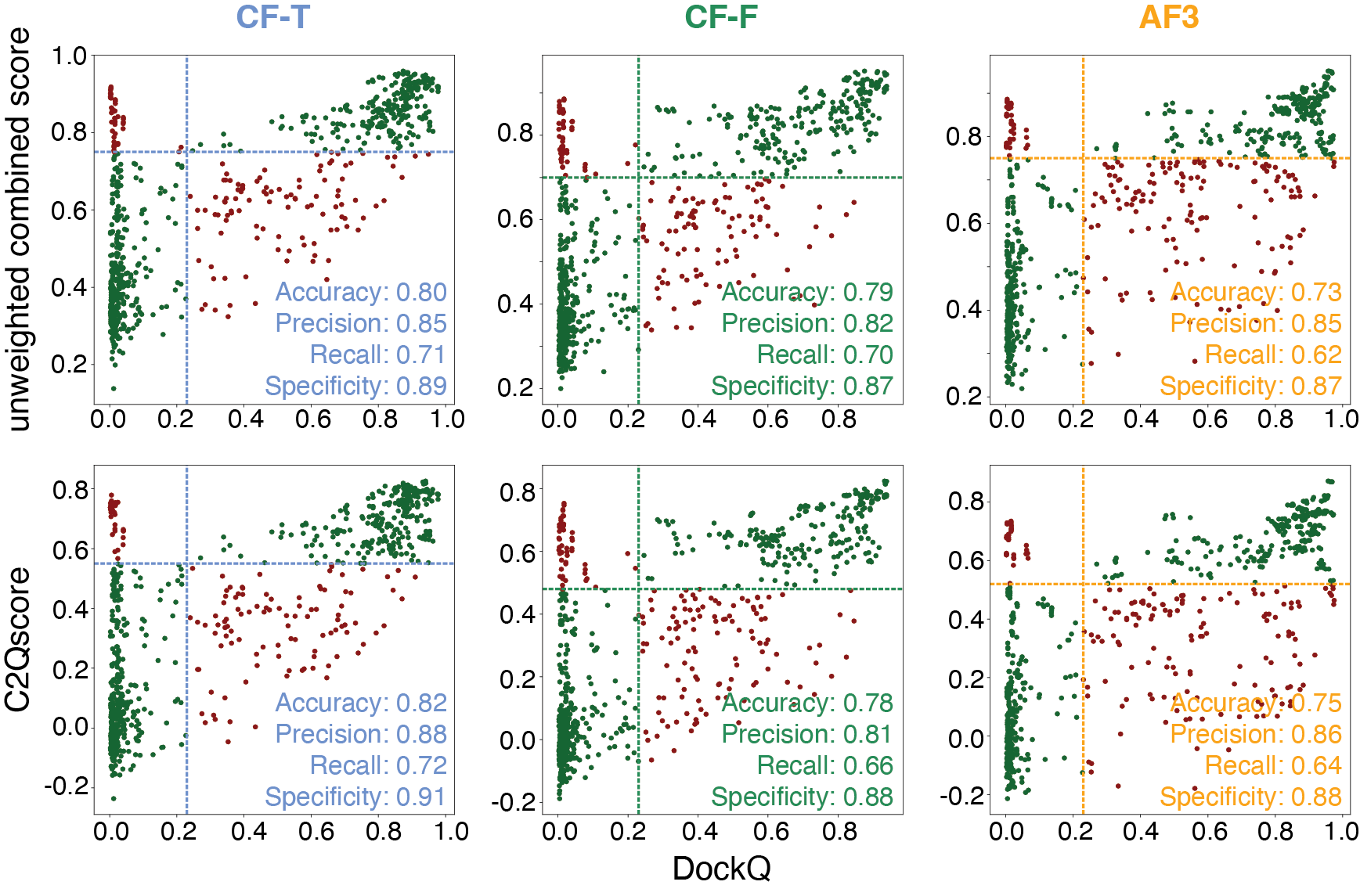

**Suppl. Fig. 8:** Unweighted combined score and C2Qscore for the large assembly test set for CF-T, CF-F and AF3. True positives and true negatives are plotted in dark green and false positives and false negatives in dark red. The cutoffs determined in Section 3.6 are shown as dotted lines in the plots. C2Qscore ranges from -0.45 to 0.88 for CF-T, -0.37 to 0.87 for CF-F and -0.33 to 0.93 for AF3.

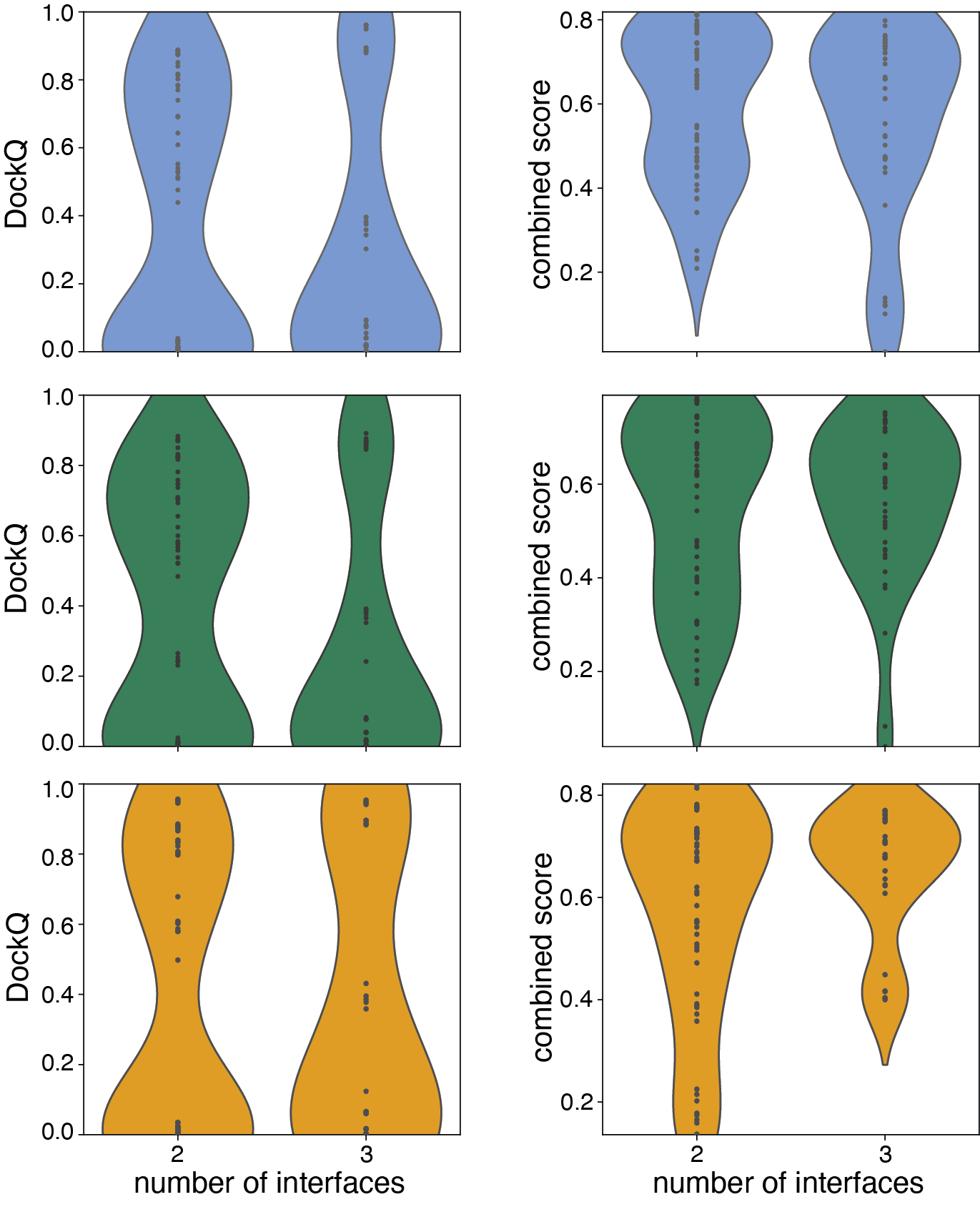

**Suppl. Fig. 9:** Distribution of DockQ and C2Qscore for CF-T cases (blue), CF-F cases (green) and AF3 cases (orange) of two (CF-T: n=58, CF-F: n=57, AF3: n=58) or three (CF-T: n=42, CF-F: n=39, AF3: n=31) potential configurations for which the prediction resembles one of the configurations.

**Suppl. Tables**

**Suppl. Tab. 1:** Prediction-based scores used for the assessment of the structural models.

| **Score** | **Range** | **What is measured?** | **Interface specific (I)/Global (G)** |
| --- | --- | --- | --- |
| pLDDT  (predicted local difference distance test)  (Jumper et al. 2021) | 0,100 | per-residue measure of the local confidence;  based on the local distance difference test Cα (Mariani et al. 2013) | G |
| PAE  (predicted alignment error)  (Evans et al. 2022) | 0, 31.75 (capped) (Varadi et al. 2022) | expected positional error [Å] at residue X, if predicted and actual structures are aligned on residue Y (confidence of the relative positions) | G |
| ipLDDT  (interface pLDDT)  (Jumper et al. 2021) | 0,100 | mean pLDDT of interface residues (residues with heavy atoms in a distance of <= 5.0Å from different chains) calculated using PICKLUSTER (Genz et al. 2023) | I |
| iPAE  (interface PAE)  (Evans et al. 2022) | 0, 31.75  (capped) (Varadi et al. 2022) | mean PAE of interface residues (residues with heavy atoms in a distance of <= 5.0Å from different chains) calculated using PICKLUSTER (Genz et al. 2023) | I |
| pTM  (Jumper et al. 2021) | 0,1 | predicted TM-score (Zhang and Skolnick 2004) | G |
| ipTM  (Evans et al. 2022) | 0,1 | measures the accuracy of the predicted relative positions of the subunits forming the protein-protein complex | I |
| *model confidence*  *(Evans et al. 2022)* | 0,1 | combination of pTM and ipTM:  0.8*ipTM+0.2*pTM | I+G |
| pDockQ2  (Zhu et al. 2023)  (https://gitlab.com/ElofssonLab/afm-benchmark) | 0,1 | predicted interface DockQ; includes an interface specific pLDDT and an interface specific PAE | I |
| VoroIF-GNN  (Olechnovič and Venclovas 2023 Jul 21)  (https://github.com/kliment-olechnovic/voronota) | 0,1 | superposition free; based on protein-protein interface graphs using Voronoi tessellation (Graphical Neural Network); estimates the accuracy of protein assemblies | I |

**Suppl. Table 2:** Accuracy and precision for individual scores for CF-T, CF-F and AF3. The used thresholds are reported in Fig. 4A.

| **Binary classification incorrect vs. (high, medium, acceptable)** | | | | | | |
| --- | --- | --- | --- | --- | --- | --- |
| **Score** | **CF-T** | | **CF-F** | | **AF3** | |
|  | **Accuracy** | **Precision** | **Accuracy** | **Precision** | **Accuracy** | **Precision** |
| ipLDDT | 0.86 | 0.91 | 0.88 | 0.93 | 0.79 | 0.92 |
| iPAE | 0.90 | 0.93 | 0.91 | 0.94 | 0.91 | 0.94 |
| pTM | 0.83 | 0.89 | 0.85 | 0.87 | 0.81 | 0.95 |
| ipTM | 0.90 | 0.93 | 0.92 | 0.94 | 0.90 | 0.96 |
| *model confidence* | 0.88 | 0.95 | 0.92 | 0.94 | 0.90 | 0.96 |
| pDockQ2 | 0.81 | 0.96 | 0.85 | 0.95 | 0.78 | 0.95 |
| VoroIF | 0.86 | 0.91 | 0.86 | 0.90 | 0.84 | 0.92 |
| **MEAN** | **0.86** | **0.93** | **0.88** | **0.92** | **0.85** | **0.94** |

**Suppl. Table 3:** Accuracy and precision for the gray zone of ipTM (0.6 <= ipTM <= 0.8) for CF-T, CF-F and AF3. The used threshold is reported in Fig. 4A.

| **Binary classification incorrect vs. (high, medium, acceptable)** | | | | | | |
| --- | --- | --- | --- | --- | --- | --- |
| **Score** | **CF-T (n = 164)** | | **CF-F (n = 180)** | | **AF3 (n = 261)** | |
|  | **Accuracy** | **Precision** | **Accuracy** | **Precision** | **Accuracy** | **Precision** |
| 0.6 <= ipTM <= 0.8 | 0.82 | 0.86 | 0.86 | 0.90 | 0.87 | 0.93 |

**Suppl. Table 4:** Pearson correlation of all approaches used to determine a combined score for CF-T, CF-F and AF3.

| **Approach** | **CF-T** | **CF-F** | **AF3** |
| --- | --- | --- | --- |
| unweighted - all scores | 0.817 | 0.829 | 0.771 |
| unweighted - Pearson correlation based | 0.820 | 0.841 | 0.779 |
| linear regression - all scores | 0.824 | 0.844 | 0.783 |
| linear regression - Pearson correlation based | 0.820 | 0.843 | 0.781 |
| lasso regression | 0.815. | 0.837 | 0.776 |

**Suppl. Table 5:** Weights for C2Qscore for CF-T, CF-F and AF3.

| **Score** | **CF-T** | **CF-F** | **AF3** |
| --- | --- | --- | --- |
| ipLDDT | 0.561 | 0.446 | -0.036 |
| iPAE | 0.123 | 0.082 | 0.169 |
| pTM | -0.084 | -0.084 | 0.335 |
| ipTM | 0.583 | 0.752 | 0.683 |
| VoroIF | 0.141 | 0.039 | 0.114 |

**Suppl. Table 6:** Bias for the unweighted combined score and C2Qscore for CF-T, CF-F and AF3

| **Approach** | **CF-T** | **CF-F** | **AF3** |
| --- | --- | --- | --- |
| unweighted - all scores | - | - | - |
| linear regression - all scores (C2Qscore) | -0.448 | -0.367 | -0.331 |

**Suppl. Table 7:** Maximal values for the unweighted combined score and C2Qscore for CF-T, CF-F and AF3

| **Approach** | **CF-T** | **CF-F** | **AF3** |
| --- | --- | --- | --- |
| unweighted - all scores | 1.000 | 1.000 | 1.000 |
| linear regression - all scores (C2Qscore) | 0.878 | 0.867 | 0.933 |

**Suppl. Table 8:** P-values for the Wilcoxon signed-rank test of C2Qscore against the unweighted combination of all scores

|  | **C2Qscore (CF-T)** | **C2Qscore (CF-F)** | **C2Qscore (AF3)** |
| --- | --- | --- | --- |
| **unweighted - CF-T** | 5.88e-184 | - | - |
| **unweighted - CF-F** | - | 5.90e-184 | - |
| **unweighted - AF3** | - | - | 5.88e-184 |

**Suppl. Table 9:** Determined thresholds for the unweighted combined score and C2Qscore for distinguishing models of low (incorrect DockQ) and high (high, medium, acceptable DockQ) quality for CF-T, CF-F and AF3

| **Approach** | **CF-T** | **CF-F** | **AF3** |
| --- | --- | --- | --- |
| unweighted - all scores | 0.75 | 0.70 | 0.75 |
| linear regression - all scores (C2Qscore) | 0.55 | 0.48 | 0.52 |
